# The HmrABCX pathway regulates the transition between motile and sessile lifestyles in *Caulobacter crescentus* by a HfiA-independent mechanism

**DOI:** 10.1101/2023.12.13.571505

**Authors:** Sébastien Zappa, Cecile Berne, Robert I. Morton, Jonathan De Stercke, Yves V. Brun

## Abstract

Through its cell cycle, the bacterium *Caulobacter crescentus* switches from a motile, free-living state, to a sessile surface-attached cell. During this coordinated process, cells undergo irreversible morphological changes, such as shedding of their polar flagellum and synthesis of an adhesive holdfast at the same pole. In this work, we used genetic screens to identify genes involved in the regulation of the motile to sessile lifestyle transition. We identified a predicted hybrid histidine kinase that inhibits biofilm formation and activates the motile lifestyle: HmrA (<u>H</u>oldfast and <u>m</u>otility regulator A). Genetic screens and genomic localization led to the identification of additional genes that regulate the proportion of cells harboring an active flagellum or a holdfast and that form a putative phosphorelay pathway with HmrA. Further genetic analysis indicates that the Hmr pathway is independent of the holdfast synthesis regulator HfiA and may impact c-di-GMP synthesis through the diguanylate cyclase DgcB pathway. Finally, we provide evidence that the Hmr pathway is involved in the regulation of sessile-to-motile lifestyle as a function of environmental stresses, namely excess copper and non-optimal temperatures.

**IMPORTANCE:** Complex communities attached to a surface, or biofilms, represent the major lifestyle of bacteria in the environment. Such a sessile state enables its inhabitants to be more resistant to adverse environmental conditions. Thus, having a deeper understanding of the underlying mechanisms that regulate the transition between the motile and the sessile states could help design strategies to improve biofilms when they are beneficial or impede them when they are detrimental. For *C. crescentus* motile cells, the transition to the sessile lifestyle is irreversible, and this decision is regulated at several levels. In this work, we describe a putative phosphorelay that promotes the motile lifestyle and inhibits biofilm formation, providing new insights into the control of adhesin production that leads to the formation of biofilms.

## INTRODUCTION

In the environment, bacteria live primarily as surface-associated colonies (1–3). Biofilms typically provide strategic advantages to their inhabitants. Indeed, cells in a biofilm can acquire transmissible DNA more easily and can be more resistant not only to xenobiotic compounds, but also to grazing by some planktonic feeding predators, phagocytosis by the host immune system, and changes in environmental conditions (4). While most studied biofilm-forming bacteria rely on a complex extracellular matrix composed of proteins, polysaccharides, DNA and other macromolecules to tightly adhere to the surface (5), many Alphaproteobacteria use a strong polar adhesin to irreversibly attach to surfaces and form biofilms (6, 7). The most extensively studied example of this type of adhesin is the *Caulobacter crescentus* holdfast.

*C. crescentus* cells have a dimorphic lifecycle, where each division cycle yields a sessile mother stalked cell and a motile daughter swarmer cell (Fig. 1A). Swarmer cells display a flagellum and multiple pili at the new pole. During the transition to the sessile form, the flagellum is ejected and the pili retracted. After losing their flagellum, cells enter the sessile phase of their life cycle. At the pole previously bearing the flagellum and pili, cells first synthesize a holdfast, followed by a cylindrical extension of the cell envelope called the stalk, which pushes the holdfast away from the cell body. The resulting stalked cells are attached to surfaces to form biofilms. Stalked cells eventually elongate, become predivisional cells, and synthesize a flagellum at the pole opposite the stalk. After cytokinesis, each newborn swarmer cell disperses and must transition to a stalked cell prior to the initiation of DNA replication and a new round of cell division.

**Figure 1:**
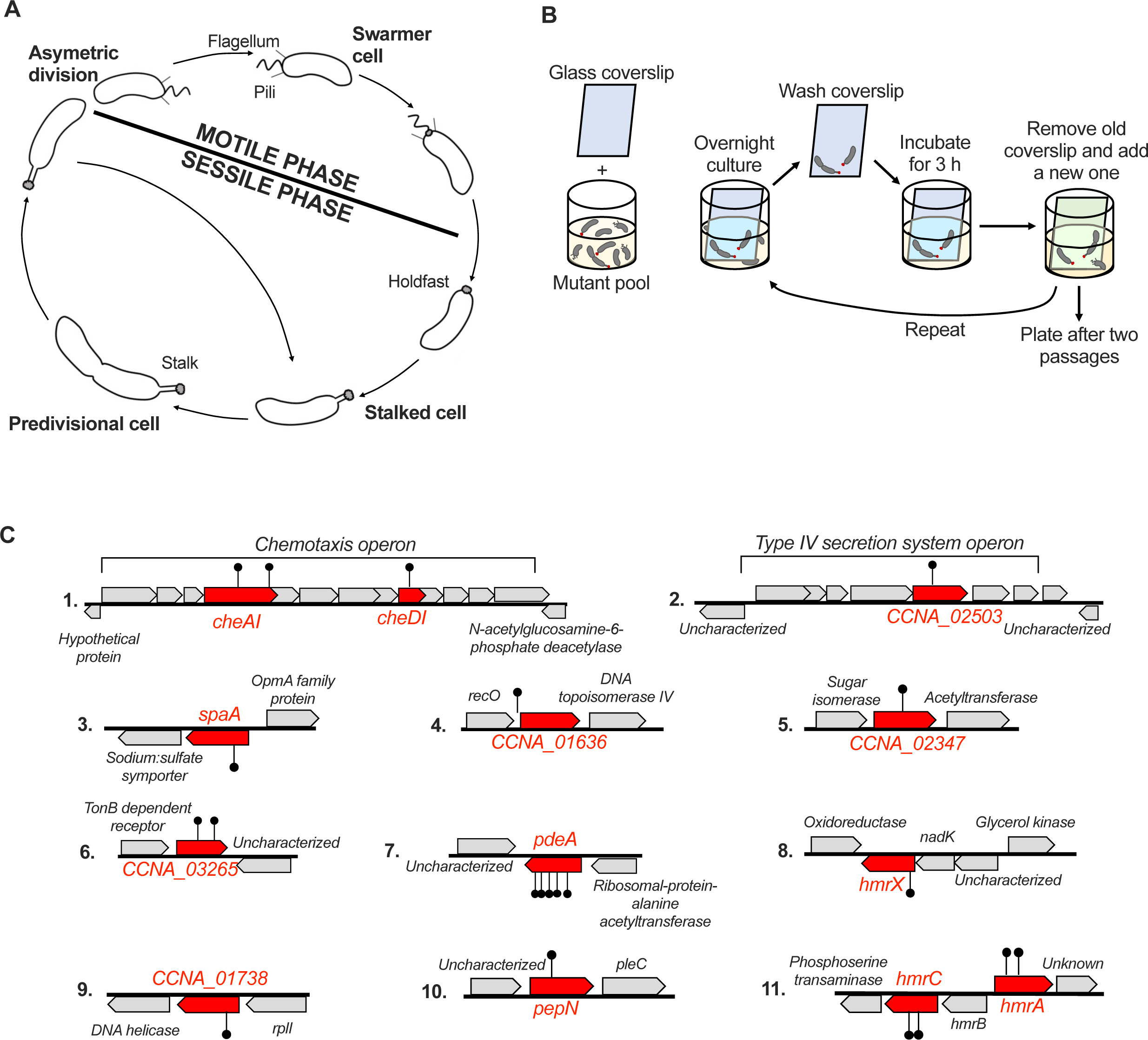
adhesion enrichment screen for identification of novel hyperadhesive mutants. **(A)** Asymmetric cell cycle of *Caulobacter crescentus.* The newborn swarmer cell harbors pili and a flagellum at the new pole of the cell. During the motile to sessile transition, this swarmer cell differentiate by retracting pili, ejecting the flagellum, and synthesizing a holdfast and a stalk at the same pole, giving rise to a stalked cell. This non-motile cell progresses through the cell cycle to give an elongated predivisional cell in which a new flagellum is synthesized at the pole opposite the stalked pole. After cell division, the newborn swarmer cell disperses and the stalked cell initiates the next round of replication. **(B)** Schematic of the forward genetic screen to identify mutants with an enhanced adhesion phenotype. A clean glass coverslip (depicted in blue) was added to a culture of pooled *Mariner* transposon mutants grown in M2X medium. After overnight growth, this coverslip was removed and thoroughly rinsed with sterile M2X, then used to inoculate a new culture. After three hours of inoculation the old coverslip (blue) was replaced with a new one (green). The enrichment process was repeated once more before isolation of single cells. Individual mutants were isolated through plating for single cells. **(C)** Identities and locations of the genes mutated in the strains isolated in the adhesion enrichment screen. Genes identified in this screen are indicated in red. The locations of the *Mariner* transposon insertions are indicated by black lollypops. The predicted functions of the proteins encoded by the flanking genes are indicated. For more information about the proteins encoded by the genes indicated in red, see Table 1.

**Table 1:**
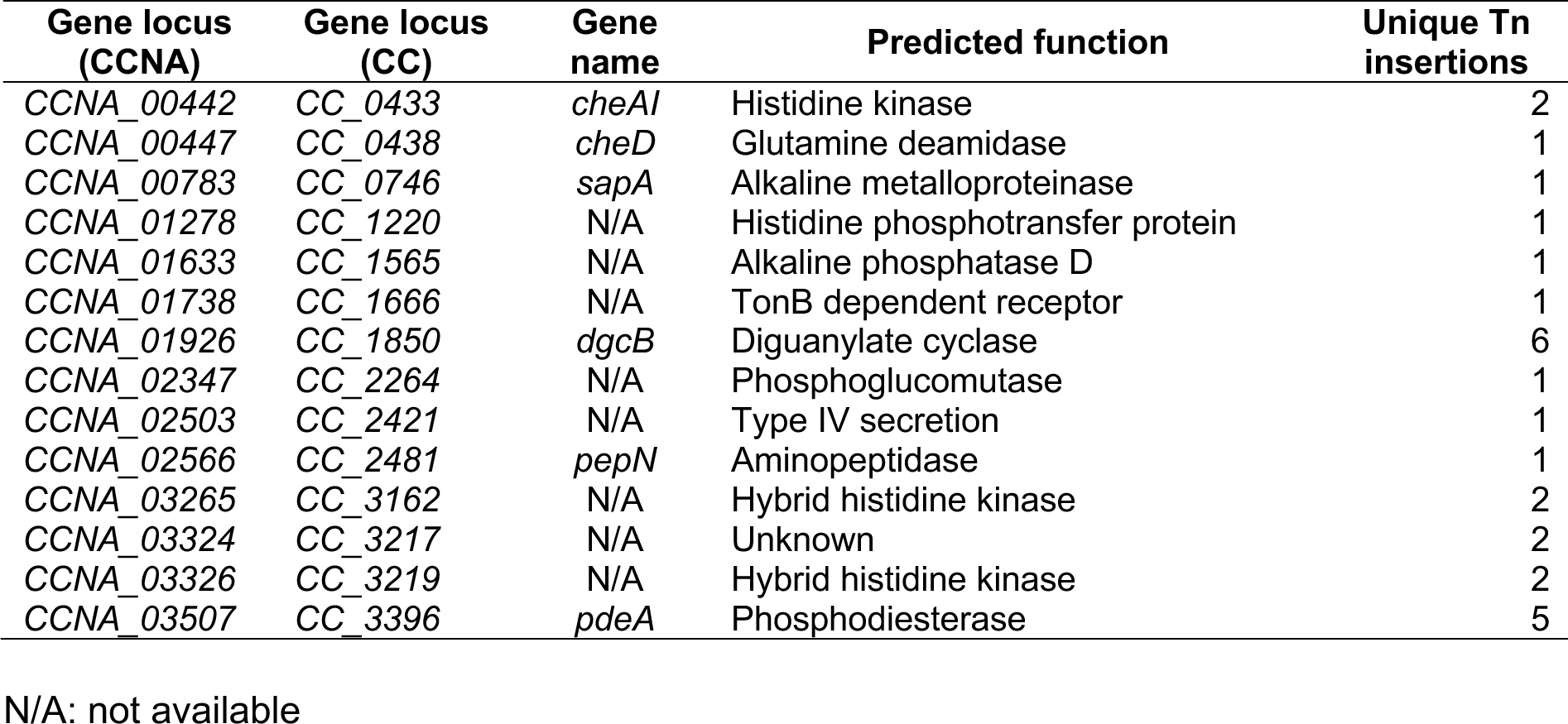
hyperadhesive mutants obtained from adhesion enrichment screen.

The progression of the cell cycle and transition between motile and sessile phases are tightly regulated. Holdfast production is temporally regulated during the cell cycle, with genes involved in holdfast synthesis under the control of key cell cycle regulators (8, 9). The intracellular concentration of cyclic di-GMP (c-di-GMP) is the primary developmental regulator of the motile to sessile lifestyle transition (10), and is involved in both flagellar ejection and holdfast production (11–13). In addition to this internal signal, holdfast production is also controlled by external environmental signals such as nutrient availability, light, and general stress responses (14–17). These signals regulate the <u>h</u>oldfast inhibitor HfiA, which inhibits the <u>h</u>oldfast <u>s</u>ynthesis protein HfsJ, a predicted glycolipid glycosyltransferase crucial for holdfast formation (15). Regulation of HfiA is controlled using different mechanisms at the transcriptional and post-translational levels, including by c-di-GMP, enabling proper timing of holdfast synthesis (15, 18, 19).

To better understand the underlying mechanisms that govern the motile to sessile transition and holdfast regulation, we performed a genetic screen for mutations that enhance biofilm formation. This work reports a putative phosphorelay that inhibits the sessile lifestyle, while promoting motility. Phosphorelays, a sub-class of two-component systems (TCS), are major regulatory pathways used by bacteria to sense and transduce environmental signals to trigger an adapted phenotypic response. A canonical TCS consists of a histidine kinase (HK) / response regulator (RR) pair. Hybrid histidine kinases (HHKs) are non-canonical TCSs, where the HK and the receiver domain of the RR are fused. HHKs represent less than 20% of bacterial HKs (20). Signal transduction from the HHK to the final RR typically involves a histidine phosphotransferase (HPT) as a mediator. Sixty-one percent of RR output domains contain a DNA-binding domain and regulate transcription (21). In addition, some output domains can transduce signals through diguanylate cyclase or phosphodiesterase domains, which produce and degrade c-di-GMP, respectively. This example highlights how regulation through phosphotransfer and c-di-GMP can be intertwined.

In this work, we describe a putative phosphorelay centered on the <u>H</u>oldfast <u>m</u>otility regulator <u>A</u> (HmrA), a HHK encoded by the *CCNA_03326* gene, identified in a screen for hyper-adhesive mutants. We show that this gene is involved in the switch from the motile to sessile lifestyle by regulating the relative number of cells harboring a flagellum or a holdfast in a mixed population. Our work shows that this mechanism is independent of HfiA and does not alter the timing of holdfast synthesis. Our data suggest that HmrA regulation of holdfast production is linked to c-di-GMP production by DgcB. In addition, we identify two putative HPTs involved in the same pathway as HmrA, namely HmrB and HmrX. Finally, we show that holdfast production is regulated by environmental signals such as the presence of excess Cu and non-optimal temperatures. This regulatory mechanism involves the protein HmrC, which is a predicted transmembrane protein required for sensing environmental stimuli in the Hmr regulation cascade.

## RESULTS

### An enrichment for hyper-adhesive mutants identifies putative elements of phosphorelays

We sought to identify mutants with an enhanced adhesion phenotype. To sensitize our screen, cells were grown in defined M2X medium in which holdfast synthesis is significantly reduced compared to using the complex medium PYE (15, 16). We randomly mutagenized a pool of *C. crescentus* NA1000 *hfsA*^+^ wild-type (WT) using the *Mariner* transposon (22). Then, as depicted in Fig. 1B, transposon mutagenized cells were pooled and grown in the presence of a glass coverslip. Cells adhering to the coverslip were recovered and enriched for two cycles. The adherent cell lines were isolated and the location of transposon insertions was determined. The enrichment resulted in 21 unique *Mariner* insertions that were mapped to 13 different genes (Table 1 and Fig. 1C). One third of the unique insertions were in genes encoding components of putative phosphorelays. Indeed, out of 21 hits, four occurred in putative HHK genes (*CCNA_03265* and *CCNA_03326*), one was in a putative HPT gene (*CCNA_01278*), and two in a gene of unknown function (*CCNA_03324*) located in the same gene cluster as the HHK *CCNA_03326*. We decided to focus on *CCNA_03326* and *CCNA_03324* since their presence in adjacent and divergent operons suggested that they might be involved in related functions. As described below, *CCNA_01278*, *CCNA_03324*, *CCNA_03325*, and *CCNA_03326* encode proteins involved in the regulation of holdfast synthesis and motility. We named these genes *hmrX*, *hmrC*, *hmrB*, and *hmrA*, respectively (for <u>h</u>oldfast and <u>m</u>otility regulators X, C, B and A).

A mutant strain harboring an in-frame deletion of the *hmrA* gene was generated to eliminate the potential polar effect of transposon insertion. The Δ*hmrA* mutant exhibited a hyper-adhesive phenotype, with approximately 7-fold more biofilm formed than the WT strain after 24h of incubation (Fig. 2A). This indicates that the putative HHK encoded by *hmrA* inhibits biofilm formation. We also constructed in-frame deletion of the other ORFs in the *hmr* gene cluster (Fig. 1C and 2A) and these mutants were tested for their ability to form biofilms. The behavior of both Δ*hmrC* and Δ*hmrB* was very similar to that of Δ*hmrA* (Fig. 2A). In contrast, Δ*03327* phenocopied the WT (Fig. 2A). We complemented each mutant strain by inserting a single copy of the deleted gene with its native promoter at the Tn*7 att* site, resulting in near WT biofilm levels for each of the complemented strains (Fig. 2A).

**Figure 2:**
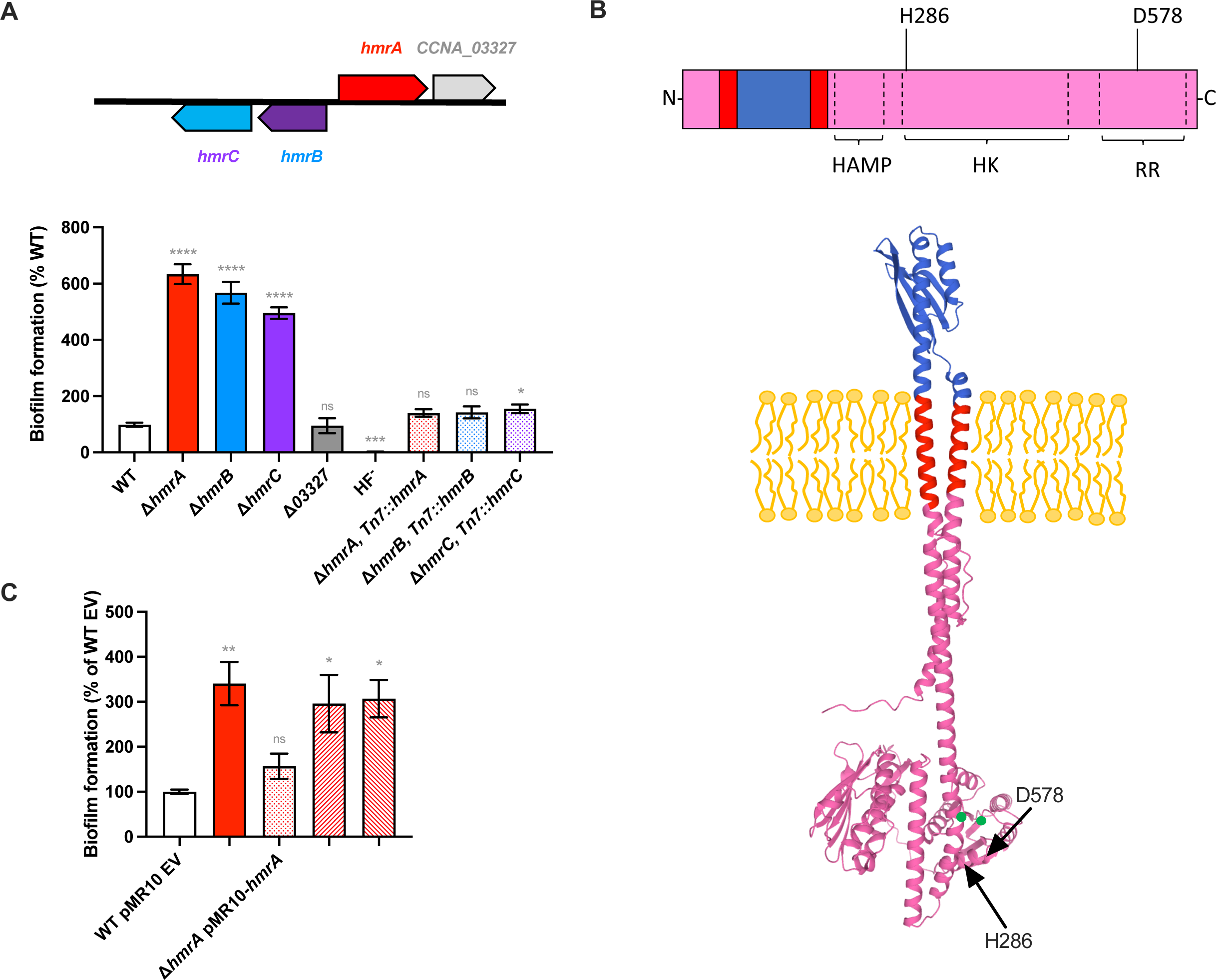
the putative HHK mutant Δ*hmrA* displays a hyperbiofilm phenotype. **(A)** Schematics of the *CCNA_03324-CCNA_03327* gene cluster (top) and biofilm formation (bottom) in 24-multiwell plates, after 24 h of incubation in M2X medium. Results are expressed as a percentage of biofilm formed by each strain compared to WT set at 100%. A holdfast-minus mutant HF^-^ (NA1000 *hfsA*^+^ Δ*hfsDAB*) was used as a no-adhesion negative control. Results are given as the average of at least three independent experiments, each run in triplicate, and the error bars represent the Standard Error of the Mean (SEM). Statistical comparisons to WT are calculated using unpaired *t*-tests (ns: not significant; * *P* < 0.05; *** *P* < 0.001; **** *P* < 0.0001). **(B)** Predicted features of HmrA structure. Top: The various predicted domains are depicted as follow: red, transmembrane helices; pink, cytoplasmic domains; blue, periplasmic domain. The putative essential H and D essential residues are indicated in position 286 and 578, respectively, as well as annotated HAMP, HK (histidine kinase) and RR (response regulator) domains. Bottom: Prediction of HmrA structure, with cellular localization, according to MembraneFold. Residues are colored according to their predicted cellular localization: red, transmembrane; pink, cytoplasmic; blue, periplasmic. **(C)** Biofilm formation in 24-multiwell plates, after 24 h of incubation in M2X medium. Results are expressed as a percentage of biofilm formed by each strain compared to WT bearing the empty pMR10 vector set at 100%. Results are given as the average of at least three independent experiments, each run in triplicate, and the error bars represent the SEM. Statistical comparisons to WT are calculated using unpaired *t*-tests (ns: not significant; * *P* < 0.05 ;** *P* < 0.005).

### HmrA is a HHK involved in biofilm regulation

The amino acid sequence of HmrA was analyzed using the conserved domain architecture database (CDART) and ExPasy Prosite (23, 24). HmrA displays typical HK and RR domains (Fig. 2B). Moreover, a HAMP linker domain was identified between amino acids (aa) 201 and 254. Such a domain is typically found in transmembrane histidine kinases where it facilitates signal transduction between a periplasmic domain and the HK domain (25). In addition, two transmembrane helices flanking a periplasmic domain, between residues 71 and 177, were predicted using the MembraneFold server for protein structure and topology prediction (26) (Fig. 2B). Thus, HmrA is likely to be a transmembrane HHK that can transduce a signal from the periplasm to the cytoplasm. The Molecular Signal Transduction database 3.0 (MiST 3.0) (27) reports 28 HHKs in the *C. crescentus* NA1000 genome. Alignment of the HmrA sequence with all 28 of these HHKs facilitated identification of the putative catalytic histidine and aspartate residues (Fig. 2B and S1A). The HK and RR domains are generally well-conserved amongst these 28 HHKs, with H286 being the best candidate for the catalytic histidine and D578 for the catalytic aspartate in HmrA. The Δ*hmrA* mutant was transformed with a low-copy replicating plasmid bearing various alleles of *hmrA*, either wild-type or harboring alanine substitutions at H286 and D578. Expression of the original *hmrA* gene in the Δ*hmrA* background reverted the phenotype to WT, while *hmrA* with H286A or D578A substitutions did not complement (Fig. 2C). In addition, a previous study showed that HmrA exhibits *in-vitro* kinase activity (28). Taken together, these data strongly suggest that HmrA acts as a HHK.

### HmrA regulates holdfast synthesis

To better assess the hyper-biofilm phenotype of the Δ*hmrA* mutant, we monitored adhesion of both WT and Δ*hmrA* strains over time. For the first few hours, WT and Δ*hmrA* strains adhered to the surface in a similar manner (Fig. 3A). However, after 5 hours post inoculation, Δ*hmrA* cells started to exhibit an enhanced adhesion phenotype that continued to increase over time (Fig. 3A). In parallel, we monitored initial adhesion events at the single cell level by recording the binding of synchronized swarmer cells to a glass surface over time by microscopy (Fig. 3B). We found that the percentage of attached cells increased at the same rate for the WT and Δ*hmrA* strains (Fig. 3B). This showed that the ability of single motile cells to interact with the surface is not impacted in the Δ*hmrA* mutant and, as a consequence, the observed enhanced biofilm formation is not due to a higher number of single swarmer cells attaching to the surface.

**Figure 3:**
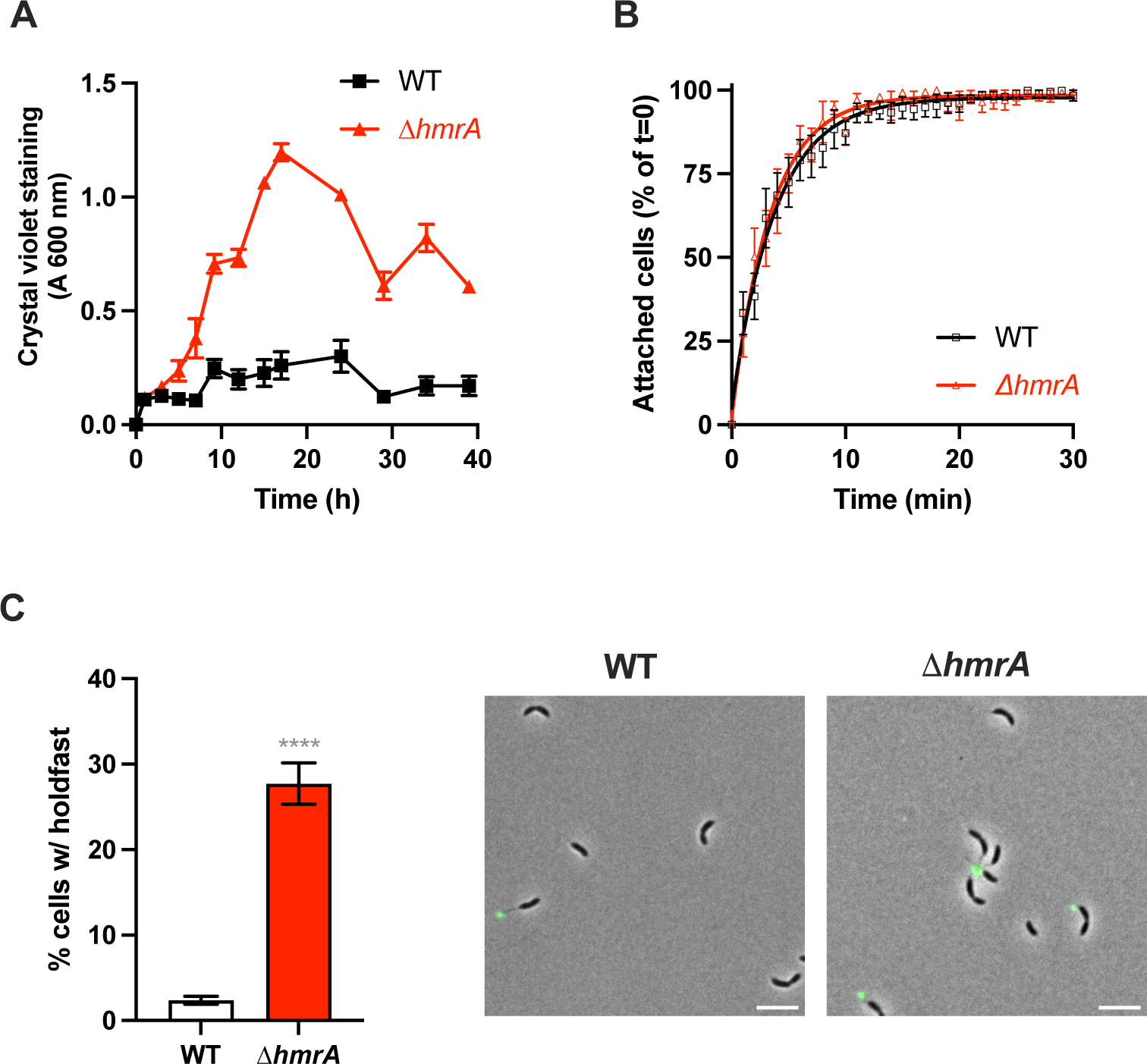
the Δ*hmrA* mutant produces more biofilm and holdfasts. **(A)** Biofilm formation over time. Amount of biofilm formed over time in M2X medium. Cultures were grown in 24-multiwell plates and the amount of biomass attached to the inside of the wells over time was quantified using crystal violet. Data are given as the average of crystal violet staining of three independent experiments, and error bars represent the SEM. **(B)** Single cell adhesion of WT (black squares) and *ΔhmrA* (red triangles) synchronized swarmer cells to a glass surface. The number of attached cells per field of view for different time points after birth is shown as a percentage of attached cells compared to the beginning of the experiment (t= 0, 0%). The attached cells are given as a percentage of the overall cell population at various times and results are averages from five samples and three independent experiments. Data were fitted using an exponential curve and error bars represent the SEM. **(C)** Left: Quantification of cells harboring a holdfast in mixed populations. The results represent the average of at least three independent replicates (more than 300 cells per replicate) and the error bars represent the SEM. Statistical comparisons to WT are calculated using unpaired *t*-tests (**** *P* < 0.0001). Right: Representative images of cells grown to mid-exponential phase in M2X and labeled with AF488-WGA to visualize holdfasts (Scale bar = 2 µm).

In *C. crescentus*, biofilm formation and permanent adhesion require the adhesive holdfast (7). Therefore, to test whether the enhanced adhesion phenotype of Δ*hmrA* could be due to the misregulation of holdfast synthesis, we first quantified the number of cells harboring a holdfast in mixed populations of cells for both WT and Δ*hmrA* strains (Fig. 3C). We found that approximately 2.5% of total WT cells harbored a holdfast, while this percentage reached more than 25% for the Δ*hmrA* mutant. Holdfast synthesis can be triggered within seconds upon contact with the surface, bypassing the developmental pathway (7, 29). However, it has been shown previously that holdfast production upon surface contact does not occur when cells are grown in nutrient limited medium such as the defined M2X medium (30). Therefore, since the increase in cells harboring a holdfast in a mixed Δ*hmrA* population occurs in the defined M2X medium, this phenotype is independent of surface contact stimulation of holdfast synthesis.

To investigate if HmrA is involved in the developmental pathway of holdfast synthesis, we monitored the timing of holdfast synthesis of newborn cells during the cell cycle by time-lapse microscopy. The duration of a complete cell cycle was identical for both WT and Δ*hmrA* strains (Fig. S2). Within that cell cycle, holdfast synthesis occurred at the same time, after approximately 30 min in both strains (Fig. S2C).

Taken together, these results suggest that HmrA is not involved in regulating the cell cycle timing of holdfast synthesis at the single cell level, but rather determines the percentage of swarmer cells that synthesize a holdfast during differentiation.

### The regulation of holdfast synthesis by HmrA is independent of HfiA

HfiA inhibits holdfast synthesis under stressful environmental conditions such as nutrient limitation, for example when growing in M2X medium (15). HfiA is regulated at different levels to ensure proper level of holdfast synthesis and its expression is controlled by a complex network of TCS including several HHKs (15, 19, 31, 32). Since we observed strong HmrA-mediated regulation of holdfast synthesis in M2X, we investigated whether it could be associated with HfiA regulation.

Biofilm formation assays showed an identical increase for Δ*hmrA* and Δ*hfiA* single mutants, and for the Δ*hmrA* Δ*hfiA* double mutant, compared to WT (Fig. S3A). Moreover, we quantified the number of cells harboring a holdfast in a mixed population and showed that this percentage increases from 15% in WT populations to 55% in Δ*hmrA* populations and reaches approximately 85% in both Δ*hfiA* and Δ*hmrA* Δ*hfiA* populations (Fig. S3B). These results indicate that HfiA still represses biofilm formation in a Δ*hmrA* mutant. In addition, we showed that the activity of the *hfiA* promoter P*_hfiA_* is not affected by the presence of HmrA (Fig. S3C). Altogether, these results indicate that inhibition of holdfast synthesis by HmrA is unlikely to occur through modulation of the HfiA regulatory pathway.

### HmrA affects cellular motility

In *C. crescentus*, holdfast misregulation is often linked to changes in motility (16, 30, 33), so we investigated the role of HmrA in motility. First, we monitored collective motility (at the population level) by measuring the swimming-dependent dispersal behavior of WT and Δ*hmrA* strains through semisolid agar M2X plates. Compared to WT, Δ*hmrA* displayed a greatly reduced swimming dispersal under these conditions: the dispersal diameter of a Δ*hmrA* strain was only a third of WT (Fig. 4A). This defective swimming dispersal phenotype could be reversed by complementing Δ*hmrA* with a single copy of *hmrA* under the control of its native promoter at the Tn*7 att* site (Fig. 4A). This result suggests that HmrA is an activator of swimming motility. Residues H286 and D578 in HmrA, which we identified previously as crucial for regulating biofilm formation (Fig. 2C), were equally important to promote dispersal through semisolid agar (Fig. 4B).

**Figure 4:**
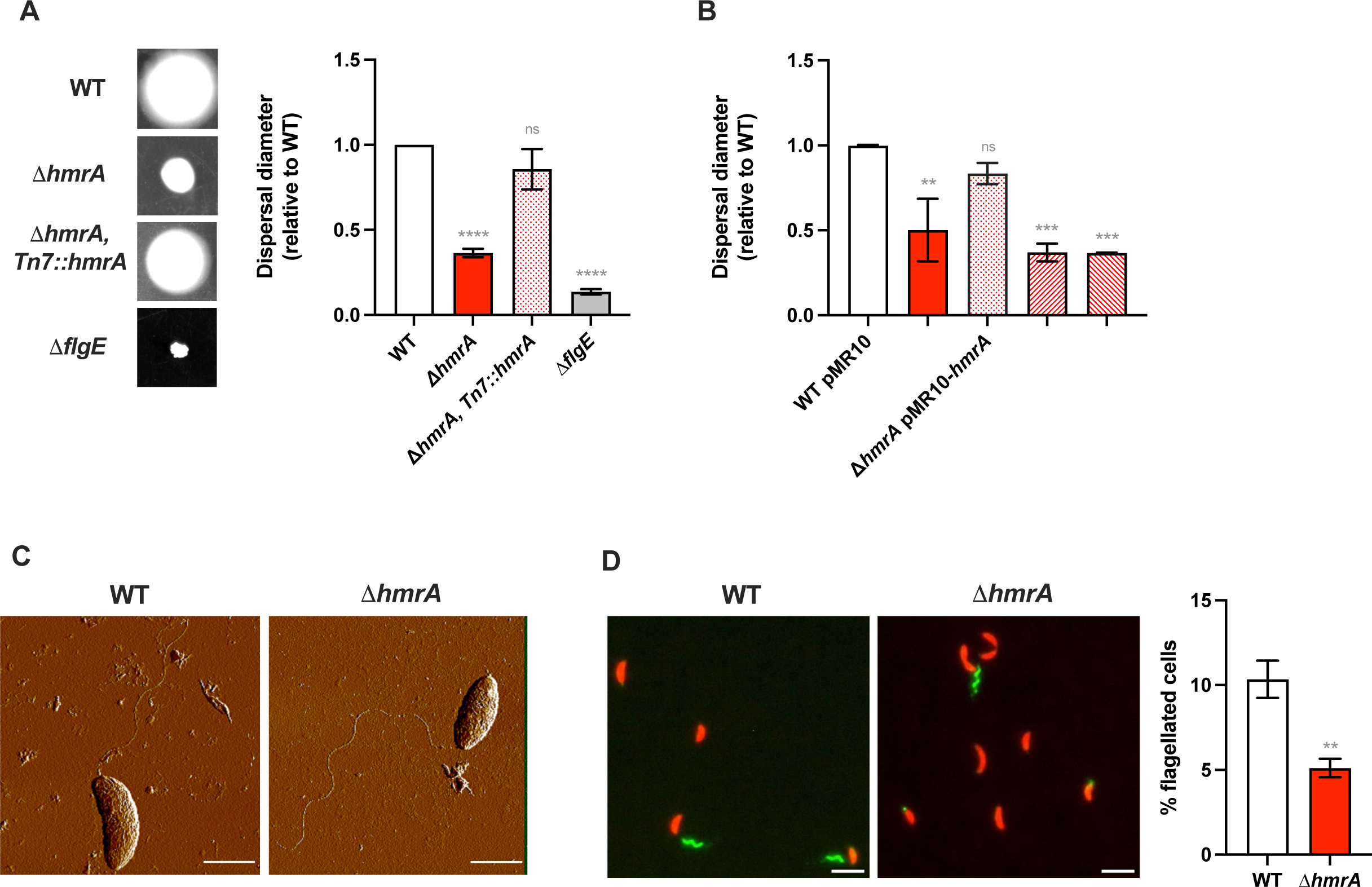
the *ΔhmrA* mutant is impaired for swimming through semisolid medium because of lower proportion of flagellated cells. **(A)** Motility assays in semisolid agar for WT and *ΔhmrA* strains, using M2X medium + 0.4% noble agar. Results are normalized to WT ring diameter measured on the same plate (set to 1). Representative images of swim rings obtained after 5 days of incubation in semisolid plates made (left). A non-motile strain lacking its flagellum (NA1000 *hfsA*^+^ Δ*flgE*) was used as a non-swimming negative control. Results represent the average of at least three independent replicates and the error bars represent the SEM. Statistical comparisons to WT are calculated using unpaired *t*-tests (ns: not significant; **** *P* < 0.0001). **(B)** Motility assays through semisolid agar for the Δ*hmrA* point mutants. Results are normalized to WT bearing the empty pMR10 vector ring diameter measured on the same plate (set to 1). Bar graphs indicate the mean of three independent replicates, and error bars represent SEM. Statistical comparisons to WT are calculated using unpaired *t*-tests (ns: not significant; * *P* < 0.05; ** *P* < 0.01; *** *P* < 0.001). **(C)** High resolution AFM images showing a WT and Δ*hmrA* swarmer cell harboring a flagellum located at the pole. Scale bar = 1 µm**. (D)** Quantification of the total number of flagella detected in mixed populations. Cells were grown to OD600 = 0.3 - 0.5 and the number of cells bearing a flagellum was scored using microscopy images where cells express mCherry (red cell bodies) and flagella are labelled using AF488-maleamide dye (green filaments). The results represent the average of at least three independent replicates (more than 500 cells per replicate) and the error bars represent the SEM. Statistical comparisons to WT are calculated using unpaired *t*-tests (** *P* < 0.005). Representative images are shown on the left (Scale bar = 2 µm).

A collective swimming defect through semisolid agar can be the consequence of various causes. We first checked if this phenotype was linked to the increased holdfast synthesis reported above. Indeed, mis-timed holdfast synthesis during the swarmer cell phase can result in a loss of motility and premature adherence to a surface (15, 34, 35). Therefore, we measured the ability of holdfast-minus (HF^-^) strains to disperse through semisolid agar. We showed that a holdfast defective NA1000 Δ*hmrA* phenotype is identical to that of a holdfast harboring NA1000 *hfaA+* Δ*hmrA* mutant (Fig. S4A). Thus, we conclude that the observed impaired swimming dispersal phenotype of the Δ*hmrA* mutant through semisolid agar is not dependent on the presence of holdfast.

We next tested whether the dispersal phenotype of the Δ*hmrA* mutant could result from a defect in chemotaxis, which can result in a decrease in the size of the dispersal ring (36, 37). In addition, it has been shown that CheA and CheB chemotaxis proteins regulate both chemotaxis and holdfast production in *C. crescentus* (16), and that the <u>C</u>heY-like c-di-GMP <u>e</u>ffector proteins CleA and CleD also play a role in regulation of holdfast synthesis (38), showing that chemotaxis, dispersal through semisolid agar, and holdfast production can be linked. To test chemotaxis, the cells were placed on a plate where xylose was introduced at the center of the plate as the sole carbon source and chemoattractant (39, 40) diffusing outward to create a gradient. Cells with a functional chemotaxis system preferentially swam towards the xylose, creating an asymmetric ring skewed towards the higher carbon concentrations. WT, Δ*hmrA* and *ΔcheAI* (a negative control that cannot perform chemotaxis (16, 41)) strains were tested, and the plates were analyzed after 48 hours (Fig. S4B). As expected, the Δ*cheAI* strain produced an evenly distributed ring, showing that it is unable to sense the xylose gradient and to display a chemotaxis phenotype. In contrast, WT and Δ*hmrA* both produced asymmetric rings towards the carbon source, showing that chemotaxis is functional in these strains. Due to its previously mentioned impaired capability in swimming dispersal, the Δ*hmrA* mutant produced a less pronounced signal than WT. Nevertheless, asymmetry of its growth ring confirmed that its chemotaxis is functional.

Finally, another cause of dispersal impairment through semisolid agar can be a defect in swimming behavior. We therefore measured the swimming speed of synchronized motile swarmer cells in liquid M2X medium and determined that single cells of WT and Δ*hmrA* display similar distributions of swimming speed (Fig. S4C). Furthermore, flagellum localization and structure were not qualitatively different, as observed by atomic force microscopy (Fig. 4C). However, when we quantified the number of cells with a flagellum in a mixed population, we found that the proportion of Δ*hmrA* cells harboring a flagellum is approximately half of WT total cells (Fig. 4D). Amongst these flagellated cells, the proportion of swarmer and predivisional cells harboring a flagellum was similar for both WT and the Δ*hmrA* strain, suggesting that the cell cycle is not perturbed (Fig. S4D).

Taken together, these results show that, while each Δ*hmrA* cell that harbors a flagellum behaves like WT, the overall proportion of flagellated cells is lower in the Δ*hmrA* population, explaining the decreased dispersal efficiency through semisolid agar. Based on these experiments, we suggest that HmrA regulates the balance between flagellum and holdfast synthesis.

### HmrA does not impact cell cycle progression

To determine if the observed differences in flagellum and holdfast formation of the Δ*hmrA* mutant is due to a misregulation of cell cycle progression, we investigated major key events in cell differentiation. First, as mentioned above, time-lapse experiments showed that the timing of the overall cell-cycle is identical in Δ*hmrA* and WT (Fig. S2). In addition, chromosome replication initiation and chromosome partitioning were similar in WT and Δ*hmrA* (Fig. S5A-C), as was the timing of swarmer to stalked cell differentiation assessed by determining the timing of PleC HK delocalization and DivJ HK localization (Fig. S5D-E). In aggregate, these results show that there is no difference in cell cycle timing and cell differentiation between WT and Δ*hmrA*.

In addition, previous studies showed that sensing of a surface by pili triggers premature cell differentiation and holdfast production in *C. crescentus* (34, 35). We therefore assessed pilus formation in Δ*hmrA* and saw no noticeable difference between WT and Δ*hmrA*. Both strains were similarly sensitive to infection with phage ΦCbK (Fig. S6A), suggesting that Δ*hmrA* produces functional pili. Moreover, the slight increase in dispersion of the Δ*pilA* mutant compared to WT is recapitulated in the Δ*hmrA* Δ*pilA* mutant compared to the Δ*hmrA* strain (Fig. S6B). Deletion of *hmrA* in the WT or Δ*pilA* background improved biofilm formation similarly by 6-8 fold compared to the parent strains (Fig. S6C). Finally, the proportion of cells bearing a holdfast in a mixed population was increased 6-7.5 times when *hmrA* was deleted in either the WT or Δ*pilA* background (Fig. S6D). These experiments strongly suggest that the Δ*hmrA* phenotype is not due to changes in pilus regulation.

Taken together, the above data show that HmrA does not control pilus formation or activity and does not regulate flagellum or holdfast formation via cell cycle timing.

### Suppressors of the Δ*hmrA* motility defect map to genes involved in c-di-GMP modulation

To gain more insight into the Hmr pathway, we performed a screen for spontaneous suppressors of the *hmrA* dispersal defect after noticing the frequent appearance of spontaneous suppressor flares through semisolid agar (Fig. 5A). Eleven of these flares (SP1 to SP11) were selected for further characterization and were tested for dispersal ability through semisolid agar and for biofilm formation (Fig. 5B and C). Their phenotypes were very similar to WT, with an increased dispersal through semisolid agar and decreased biofilm formation compared to the Δ*hmrA* parent strain. Only the SP9 suppressor displayed a level of biofilm formation that was similar to its parental strain Δ*hmrA*, while its dispersal through semisolid agar was similar to WT, like the rest of the suppressors. The location and nature of the different mutations are summarized in Table 2 and Fig. 5D. Eight out of 11 spontaneous suppressors had distinct mutations in the diguanylate cyclase gene *dgcB* (*CCNA_01926*) (Table 2 and Fig. 5D), a known regulator of holdfast production (13). DgcB is involved in the mechanosensing pathway of holdfast production through interaction with the flagellar motor to trigger production of c-di-GMP, leading to the c-di-GMP-dependent activation of the holdfast synthesis protein HfsJ (42, 43).

**Figure 5:**
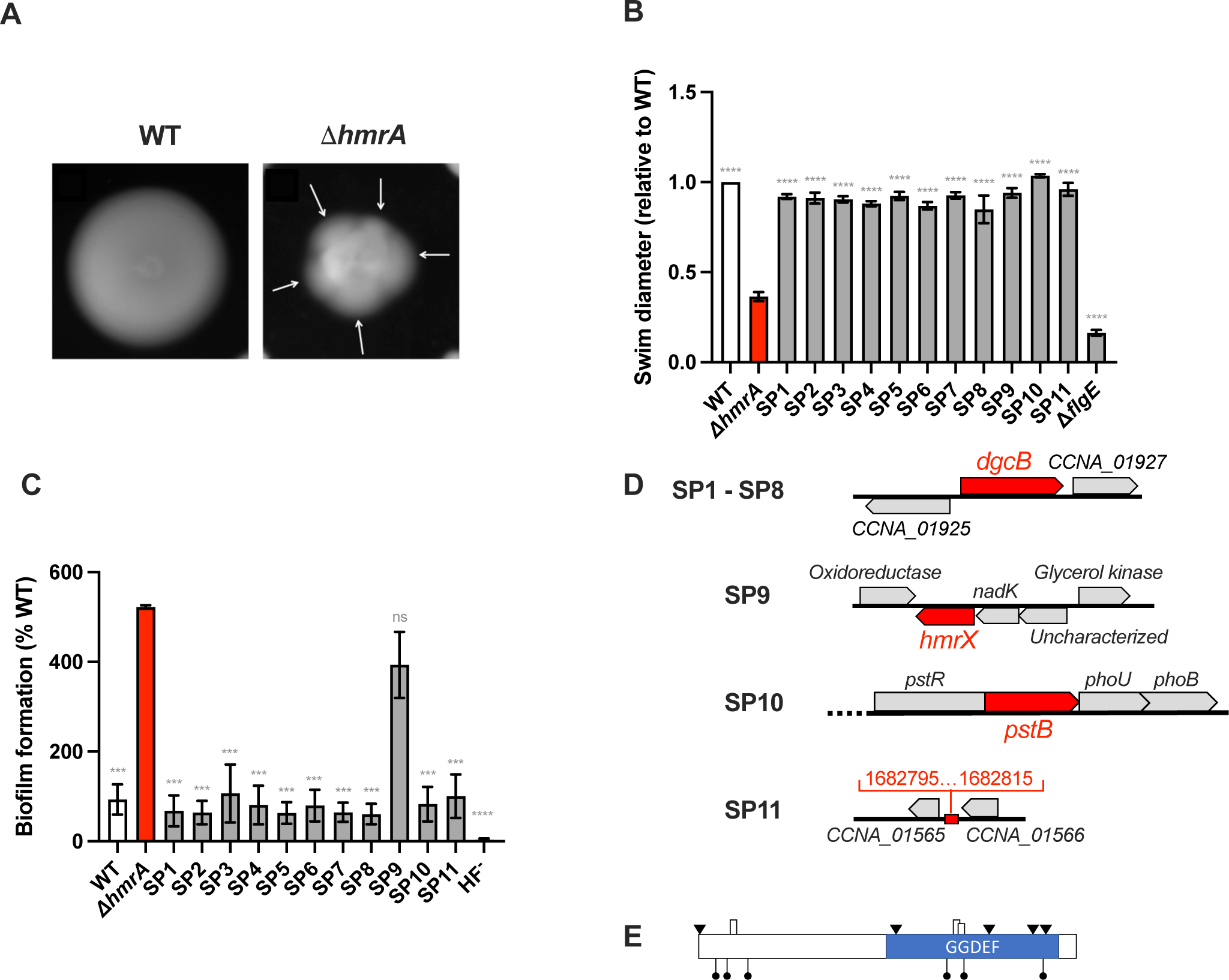
identification of mutants that suppress the motility deficiency of Δ*hmrA* in semisolid agar plates. **(A)** Representative examples of semisolid agar plates inoculated with WT (left) and the Δ*hmrA* mutant (right). Both strains were inoculated into a PYE agar plate containing 0.4% noble agar and imaged after 7 days at room temperature. The arrows indicate flares of spontaneous suppressors in the Δ*hmrA* background. **(B)** Motility assays through semisolid agar. Results are normalized to WT ring diameter measured on the same plate (set to 1). Bar graphs indicate the mean of three independent replicates, and error bars represent SEM. Statistical comparisons to Δ*hmrA* are calculated using unpaired *t*-tests (**** *P* < 0.0001). **(C)** Biofilm formation of Δ*hmrA* suppressors. Results are expressed as a percentage of biofilm formed by each strain compared to WT set at 100%. Results are given as the average of three independent experiments, each run in triplicate, and the error bars represent the SEM. Statistical comparisons to Δ*hmrA* are calculated using unpaired *t*-tests (ns: not significant; *** *P* < 0.001; **** *P* < 0.0001). **(D)** Identities and locations of the genes mutated in the strains isolated in the adhesion enrichment screen. Genes that were identified are indicated in red. For more information about these mutations, see Table 2. **(E)** Schematics of the *dgcB* gene and location of the mutations found in the different suppressor strains (SP1 to SP8). The locations of the *Mariner* transposon insertions are indicated by black lollipops, point mutations are illustrated as black triangles and insertions/deletions as boxes. For more information about these mutations, see Table 2.

**Table 2:** Δ*hmrA* suppressors displaying semisolid swimming phenotype.

| Suppressor name | Gene locus (CCNA) | Gene locus (CC) | Gene name | Mutation <sup>a</sup> | Frameshift mutation |
| --- | --- | --- | --- | --- | --- |
| SP1 | CCNA_01926 | CC_1850 | *dgcB* | ΔG96- C115 | Yes |
| SP2 | CCNA_01926 | CC_1850 | *dgcB* | ΔT740- G757 | No |
| SP3 | CCNA_01926 | CC_1850 | *dgcB* | ΔC751- C768 | No |
| SP4 | CCNA_01926 | CC_1850 | *dgcB* | A573T | No |
| SP5 | CCNA_01926 | CC_1850 | *dgcB* | G827::<sup>*</sup> <sup>b</sup> | Yes |
| SP6 | CCNA_01926 | CC_1850 | *dgcB* | T955C | No |
| SP7 | CCNA_01926 | CC_1850 | *dgcB* | A986::*A* | Yes |
| SP8 | CCNA_01926 | CC_1850 | *dgcB* | T1::*A* | Yes |
| SP9 | CCNA_01278 | CC_1220 | N/A | A152T | No |
| SP10 | CCNA_00294 | CC_0292 | *pstB* | T275P | No |
| SP11 |  |  |  | A1682805T <sup>c</sup> | No |

<sup>a</sup> Mutations indicated in the mutation column were identified through sequencing of *dgcB* or the entire genome.
<sup>b</sup> The \* in SP5 represents the insertion of the following DNA sequence ACCACGCTGGAGGAAACG CGAAGCCGCCTCGGTGGTGGCG.
<sup>c</sup> The mutation in SP11 mutant is in an intergenic region and the numeric site of the mutation is indicated.

In parallel, we also performed a similar suppressor screen by *Mariner* transposon mutagenesis and identified transposon insertions that restored dispersal to Δ*hmrA* through semisolid agar plates. All of the 6 analyzed mutants presented unique insertions in *dgcB* (Fig. 5E). Interestingly, *dgcB* was the only diguanylate cyclase gene identified in both Δ*hmrA* motility suppressor screens. It has been shown that the diguanylate cyclases PleD and DgcB are involved in flagellum and holdfast regulation via the developmental and mechanical pathway, respectively (42). Therefore, we constructed *dgcB* or *pleD* mutants in the Δ*hmrA* background to determine if HmrA is involved in the mechanical or developmental holdfast production pathway, or both (Fig. 6A and B). Figure 6B shows that deleting *dgcB* or *pleD* in a WT background decreased the number of cells harboring a holdfast by approximately 40% (2.9 ± 0.8, 1.8 ± 1.6, and 1.7 ± 1.5 % of number of cells harboring a holdfast for WT, Δ*dgcB,* and *ΔpleD* respectively). In contrast, deleting *hmrA* in a WT or Δ*pleD* background caused an increase in cells bearing a holdfast in the population by 8-10 times (22.6 ± 11 and 16.5 ± 2.2 for *ΔhmrA* and *ΔhmrA ΔpleD* respectively). The Δ*hmrA* Δ*dgcB* mutant only produces around 3 times more holdfasts (5.2 ± 1.6 %) than the Δ*dgcB* single mutant. Thus, while deleting *hmrA* in a *pleD* background phenocopies a *hmrA* deletion in WT, this is not the case in a *dgcB* background. These results strongly suggest that the *hmrA* phenotypes are specifically due to an increase in DgcB-dependent c-di-GMP synthesis and linked to the mechanical pathway of holdfast production (33). In support of this hypothesis, the aforementioned suppressor screens revealed DgcB as the sole putative diguanylate cyclase partner for HmrA.

**Figure 6:**
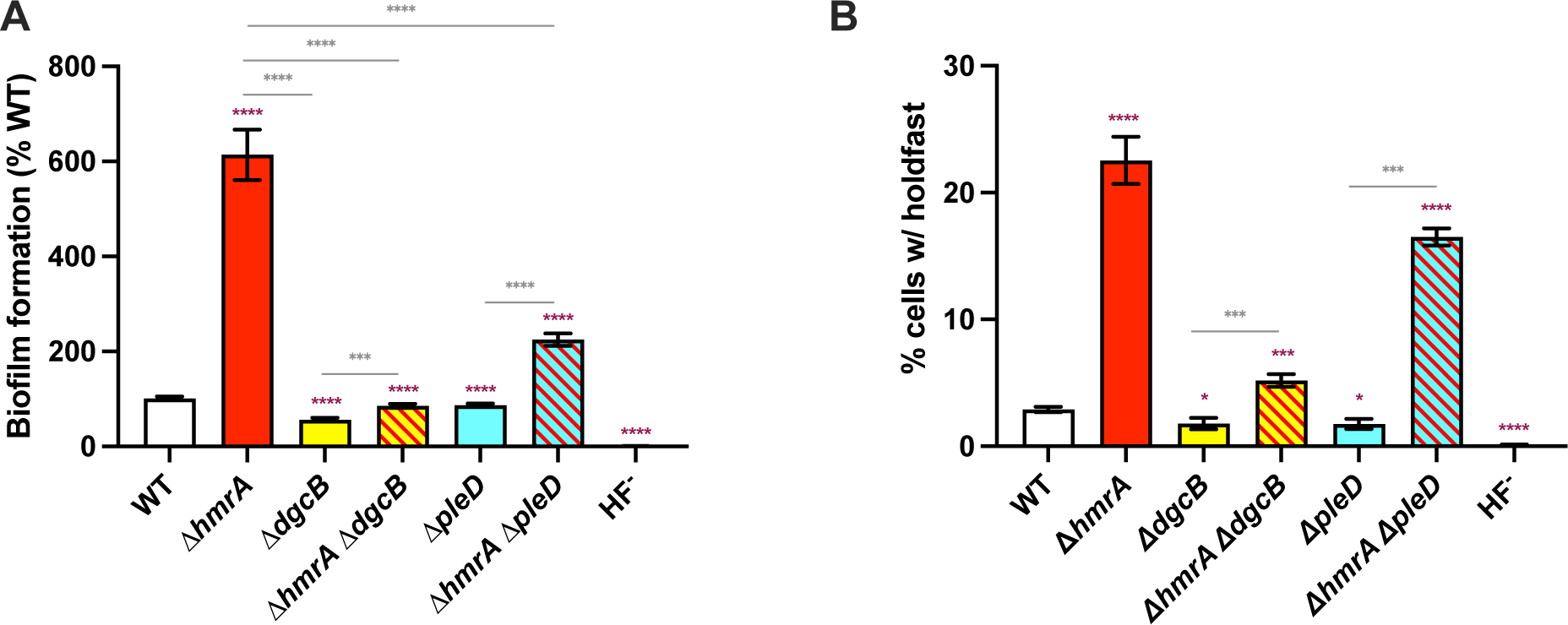
biofilm and semisolid agar swimming of the Δ*dgcB* and Δ*pleD* mutants. **(A)** Biofilm formation. Results are expressed as a percentage of biofilm formed by each strain compared to WT set at 100%. Results are given as the average of three independent experiments, each run in triplicate, and the error bars represent the SEM. **(B)** Quantification of cells harboring a holdfast in mixed populations. The results represent the average of at least three independent replicates (more than 300 cells per replicate) and the error bars represent the SEM. For both graphs, statistical comparisons are calculated using unpaired *t*-tests. Comparisons to WT are shown in maroon while other comparisons are shown in gray (* *P* < 0.05; ** *P* < 0.01; *** *P* < 0.001; **** *P* < 0.0001).

### HmrX and HmrB are putative histidine-phosphotransferases working in concert with HmrA

Another spontaneous *hmrA* suppressor mutant that caught our attention is SP9 (Table 2). This mutant has a A152T mutation in HmrX. According to the MembraneFold server for protein structure and topology prediction (26), the A152 residue is located in a four-helix bundle with access to the surrounding solvent (Fig. 7A). A four-helix bundle is a feature of HTPs, otherwise poorly conserved in terms of primary sequence. Moreover, HmrX is predicted to be a soluble cytoplasmic protein. Finally, analysis of HmrX predicted tridimensional structure using the Dali server (44) returns the chemotaxis kinase CheA and a putative histidine kinase as top hits. Overall, HmrX is predicted to be a small soluble cytoplasmic protein consisting of a four-helix bundle and with sequence similarity to proteins involved in phosphorelays. This supports the HPT nature of HmrX, as it was previously suggested from an *in silico* search for HPTs in the *C. crescentus* genome (45). To determine if HmrX is part of the HmrA phosphorelay, we constructed a strain harboring an in-frame deletion of *hmrX*, and tested its dispersal through semisolid agar, adhesion, and holdfast production capabilities (Fig. 7B-D).

**Figure 7:**
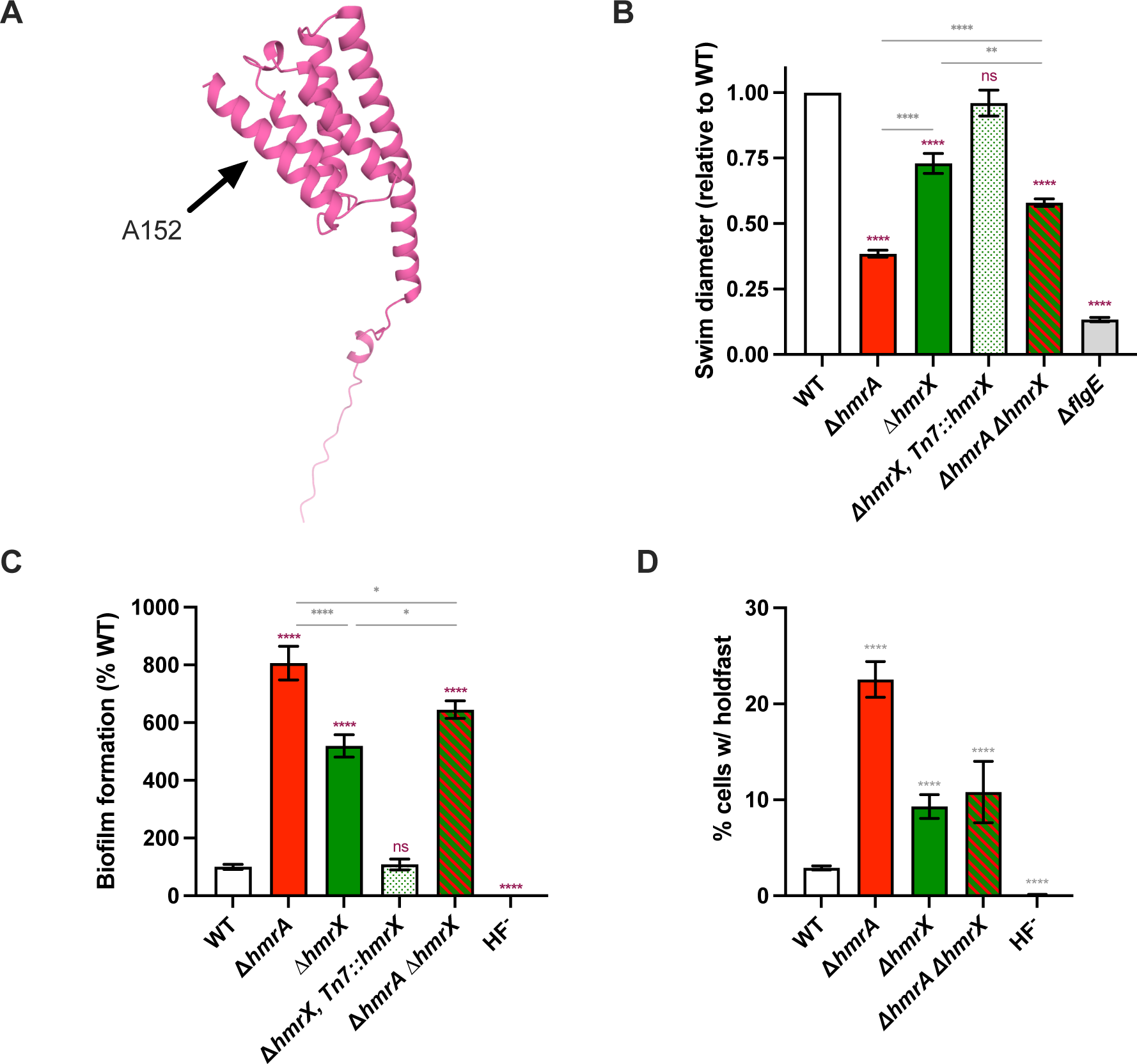
HmrX, a putative HTP involved in the Hmr pathway. **(A)** Prediction of HmrX tridimensional structure, with cellular localization, according to MembraneFold. Residues are colored according to their predicted cellular localization (pink – cytoplasmic). **(B)** Motility assays through semisolid agar for the Δ*hmrA* spontaneous mutants. Results are normalized to WT ring diameter measured on the same plate (set to 1). Bar graphs indicate the mean of three independent replicates, and error bars represent SEM. Statistical comparisons are calculated using unpaired *t*-tests. Comparisons to WT are shown in maroon while other comparisons are shown in gray (ns: not significant; ** *P* < 0.01; **** *P* < 0.0001)**. (C)** Biofilm formation. Results are expressed as a percentage of biofilm formed by each strain compared to WT set at 100%. Results are given as the average of three independent experiments, each run in triplicate, and the error bars represent the SEM. Statistical comparisons are calculated using unpaired *t*-tests. Comparisons to WT are shown in maroon while other comparisons are shown in gray (ns: not significant; * *P* < 0.05; **** *P* < 0.0001). **(D)** Quantification of cells harboring a holdfast in mixed populations. The results represent the average of at least three independent replicates (more than 300 cells per replicate) and the error bars represent the SEM. Statistical comparisons to WT are calculated using unpaired *t*-tests (**** *P* < 0.0001).

The Δ*hmrX* mutant displayed a smaller swim diameter in semisolid agar plates compared to WT (Fig. 7B). This phenotype was fully restored to WT levels when the mutant was complemented with one copy of *hmrX* (Fig. 7B). In addition, biofilm formation and number of cells bearing a holdfast were increased in the Δ*hmrX* mutant (Fig. 7C-D). In summary, deletion of *hmrX* yielded phenotypes that were similar to those of the Δ*hmrA* mutant, although their amplitude was less pronounced. Interestingly, the Δ*hmrA* Δ*hmrX* double mutant displayed swimming and biofilm phenotypes that were intermediate between the strong Δ*hmrA* phenotype and the more modest Δ*hmrX* phenotype. Regarding the ratio of cells harboring holdfast, Δ*hmrX* was epistatic to *hmrA*, suggesting that the putative HPT HmrX is part of the Hmr pathway, which is also supported by its identification as a suppressor of Δ*hmrA*.

Furthermore, although *CCNA_03325* (*hmrB*) was not identified in our genetic screens, we investigated its function since it is adjacent to *hmrA* and *hmrC* (Fig. 2A) and its protein product meets two of the three criteria previously used to search for HPT proteins in *C. crescentus* (45): it is less than 250 aa (108 aa), and is predicted to contain more than 70% of alpha helices (87%). However, the HxxKG motif, used as a third criterium, is not present in HmrB, but it is not clear if this motif is a reliable signature of HPT (46). More importantly, the typical four-helix bundle conserved in HPT is not observed in the predicted tridimensional structure of HmrB (Fig. 8A). Instead, according to MembraneFold, it consists of a two-helix bundle of 35-40 aa each, making up most of a soluble cytoplasmic protein. Analysis of this predicted structure using the Dali server did not return any strong hint to predict its function. Phenotypic analysis showed that the Δ*hmrB* mutant displays: i) a strong hyperbiofilm phenotype (Fig. 2A and 8B); ii) a high ratio of cells harboring a holdfast (Fig. 8C); and iii) impaired swimming motility through semisolid agar (Fig. 8D). These three phenotypes are comparable to those of the Δ*hmrA* mutant and can be reversed to WT levels by complementation with a single copy of *hmrB* under its native promoter (Fig. 2A and 8D). The Δ*hmrA* Δ*hmrB* double mutant behaved similarly to the Δ*hmrA* and Δ*hmrB* single mutants (Fig. 8B-D), suggesting that HmrA and HmrB act in the same pathway. Finally, assuming HmrB is an atypical HTP, its phosphotransferase activity could be achieved by one of the two histidine residues, H50 and H74. The predicted tridimensional structure indicates H50 as accessible by the solvent while H74 is located on one of the two helices and facing the other one (Fig. 8A). Thus, H50 is a better candidate as the catalytic residue for phosphotransfer. Both histidine residues were substituted with an alanine and neither of the substituted versions could rescue biofilm formation or swimming motility of the Δ*hmrB* strain (Fig. 8E-F). We suggest that substituting H50 disrupts phosphotransfer while mutating H74 disrupt structural stability (Fig. 8A). In aggregate, these findings suggest that *hmrB* codes for an atypical HPT that acts in concert with HmrA.

**Figure 8:**
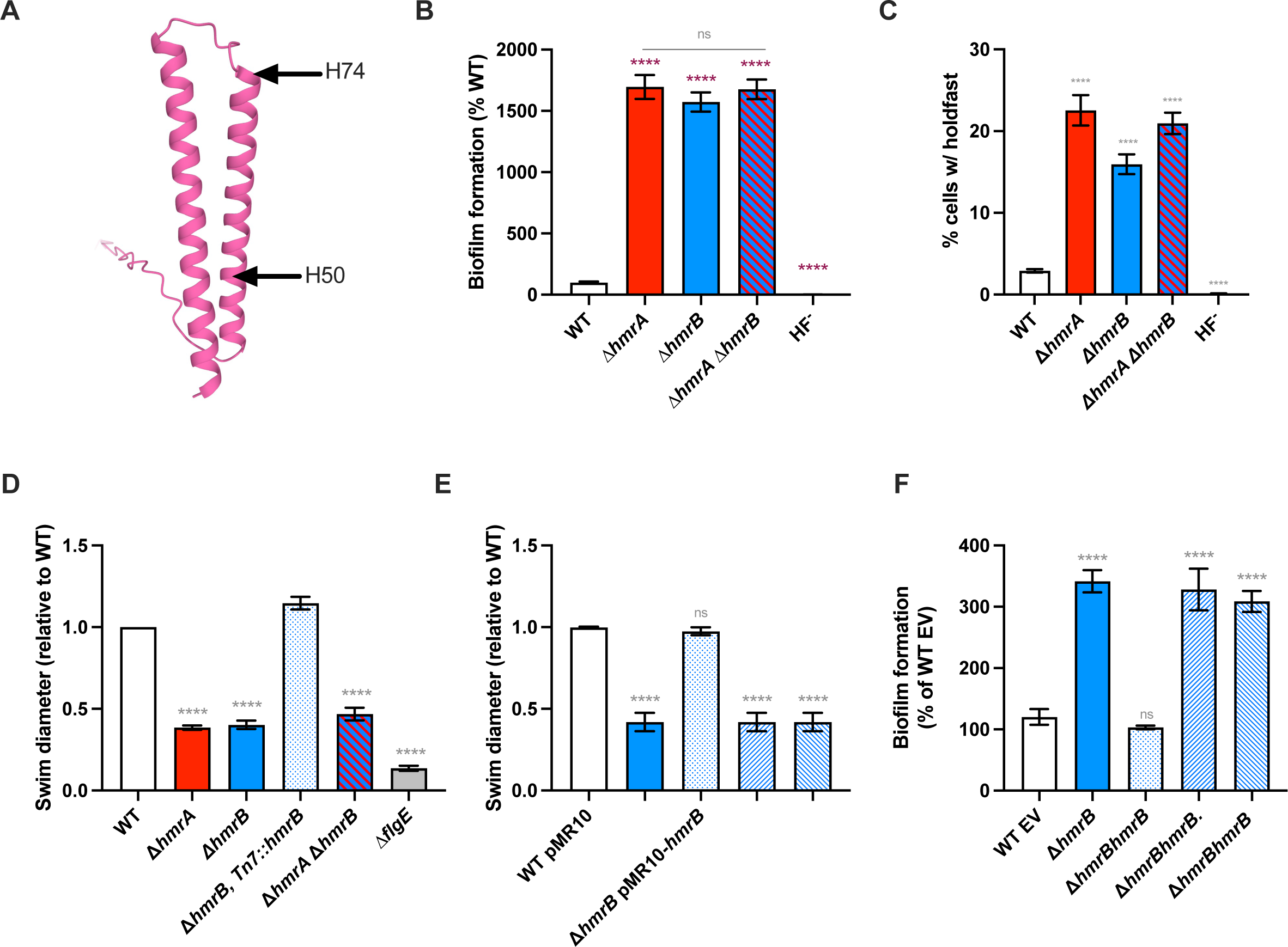
HmrB, a putative atypicial HTP involved in the Hmr pathway. **(A)** Prediction of HmrB tridimensional structure, with cellular localization, according to MembraneFold. Residues are colored according to their predicted cellular localization pink – cytoplasmic). **(B)** Biofilm formation. Results are expressed as a percentage of biofilm formed by each strain compared to WT set at 100%. Results are given as the average of three independent experiments, each run in triplicate, and the error bars represent the SEM. Statistical comparisons are calculated using unpaired *t*-tests. Comparisons to WT are shown in maroon while other comparisons are shown in gray (ns: not significant; **** *P* < 0.0001). **(C)** Quantification of cells harboring a holdfast in mixed populations. The results represent the average of at least three independent replicates (more than 300 cells per replicate) and the error bars represent the SEM. Statistical comparisons to WT are calculated using unpaired *t*-tests (**** *P* < 0.0001). **(D-E)** Motility assays through semisolid agar. Results are normalized to WT ring diameter measured on the same plate (set to 1). Bar graphs indicate the mean of three independent replicates, and error bars represent SEM. Statistical comparisons to WT are calculated using unpaired *t*-tests (**** *P* < 0.0001). **(F)** Biofilm formation. Results are expressed as a percentage of biofilm formed by each strain compared to WT set at 100%. Results are given as the average of three independent experiments, each run in triplicate, and the error bars represent the SEM. Statistical comparisons to WT are calculated using unpaired *t*-tests (ns: not significant; **** *P* < 0.0001).

### HmrC is required for sensing environmental stress and functions upstream of HmrA

Finally, we analyzed the third gene in the *hmrCBA* cluster, which was the site of two transposon insertions in our initial genetic screen, *hmrC* (Fig. 1C and Table 1). In-frame deletion of this gene yielded increased biofilm formation, a higher ratio of cells harboring a holdfast, and decreased swimming though semisolid agar, all at comparable levels to Δ*hmrA* and Δ*hmrB* (Fig. 2A and 9A-C). Performing these experiments with the Δ*hmrA* Δ*hmrC* double mutant resulted in the same phenotypes as with the single Δ*hmrA* and Δ*hmrC* mutants. From this set of experiments, we propose that HmrA and HmrC function in the same regulatory pathway.

**Figure 9:**
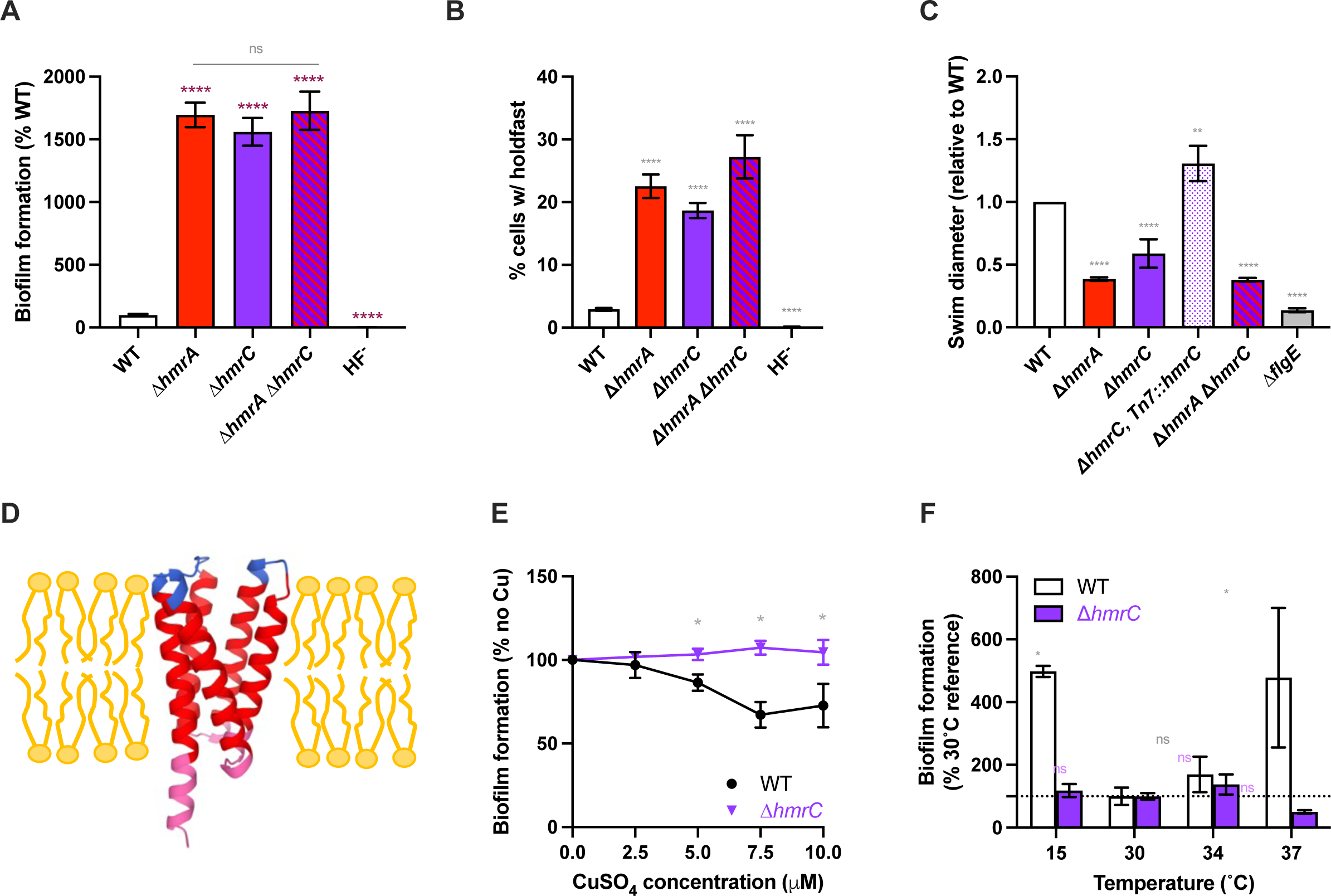
HmrC, a putative membrane protein sensing environmental stress and involved in the Hmr pathway. **(A)** Biofilm formation. Results are expressed as a percentage of biofilm formed by each strain compared to WT set at 100%. Results are given as the average of three independent experiments, each run in triplicate, and the error bars represent the SEM. Statistical comparisons are calculated using unpaired t-tests. Comparisons to WT are shown in maroon while other comparisons are shown in gray (ns: not significant; **** P < 0.0001). **(B)** Quantification of cells harboring a holdfast in mixed populations. The results represent the average of at least three independent replicates (more than 300 cells per replicate) and the error bars represent the SEM. Statistical comparisons to WT are calculated using unpaired t-tests (**** P < 0.0001). **(C)** Motility assays in semisolid agar. Results are normalized to WT ring diameter measured on the same plate (set to 1). Bar graphs indicate the mean of three independent replicates, and error bars represent SEM. Statistical comparisons to WT are calculated using unpaired t-tests (** P < 0.01; **** P < 0.0001). **(D)** Prediction of HmrC tridimensional structure, with cellular localization, according to MembraneFold. Residues are colored according to their predicted cellular localization: red - transmembrane; pink – cytoplasmic; blue: extracellular. **(E)** Biofilm formation in M2X medium adding different concentration of CuSO_4_ for WT (black circles) and Δ*hmrC (*purple down triangles). Results are expressed as a percentage of biofilm formed by each strain in the presence of metal compared to the no metal addition set at 100%. Results are given as the average of three independent experiments, each run in triplicate, and the error bars represent SEM. Statistical comparisons to the no metal added controls are calculated using unpaired t-tests (ns: not significant; ** P < 0.01; **** P < 0.0001). **(F)** Biofilm formation in M2X medium using different incubation temperatures. Results are expressed as a percentage of biofilm formed by each strain at a given temperature compared to the 30°C reference set at 100%. Results are given as the average of three independent experiments, each run in triplicate, and the error bars represent SEM. Statistical comparisons to the 30°C reference conditions are calculated using unpaired t-tests (ns: not significant; * P < 0.05).

Genome annotation did not ascribe a function to HmrC and a BLAST search did not return any conserved domain. Nevertheless, using MembraneFold, which combines protein structure prediction (AlphaFold and OmegaFold) and transmembrane topology (DeepTMHMM), analysis of the sequence showed that this 142 aa protein is predicted to contain five transmembrane helices, which make up 57% of the protein (Fig. 9D). Moreover, the I-TASSER server for prediction of protein structure and function (47) revealed that *hmrC* encodes a protein with similarity to the substrate binding component (S-component) of an energy-coupling factor (ECF) type transporter (48). Finally, analysis of the predicted tridimensional structure from AlphaFold with the Dali server returned hits involved in signal sensing/transducing and phosphotransfer.

Previous whole-genome transcriptome analyses revealed that *C. crescentus hmrC* is regulated by environmental stresses, namely metal exposure and cold shock (49, 50). Indeed, the UzcR response regulator component of the two-component system UzcRS, which is involved in the sensing of U, Zn and Cu in *C. crescentus*, regulates *hmrC* by interacting with its promoter (49). Thus, we hypothesized that the Hmr pathway plays a role in growth in the presence of Cu and Zn. We tested the ability of WT and mutant strains to make biofilms in the presence of increasing concentrations of copper (CuSO_4_) and zinc (ZnSO_4_). We identified CuSO_4_ and ZnSO_4_ concentration ranges that affect biofilm formation without impacting cell growth for subsequent testing (Fig. S7). While WT biofilm formation is impaired by increased concentration of CuSO_4_ from 5 to 10 µM, it is enhanced by addition of up to 100 µM ZnSO_4_ (Fig. 9E and S7). However, biofilm formation of the Δ*hmrC* mutant was not impacted by the addition of these metals (Fig. 9E and S7). The same trend was observed in the presence of ZnSO_4_ (Fig. S7C), but the difference is not statistically significant. This could be explained by the variability in the response to metal addition was observed during these experiments, with the amplitude of the WT response to metals strongly fluctuating between replicates. Thus, we conclude that HmrC acts as an inhibitor of biofilm formation in the presence of excess copper.

HmrC was also found to be regulated in response to cold exposure by RNA-seq (50) Therefore, we decided to monitor the biofilm formation of both WT and Δ*hmrC* at a range of temperatures, from 15°C to 37°C (Fig. 9F). We noticed that at temperatures 15°C and 37°C, WT forms around 5 times more biofilms compared to temperatures closer to the optimal growth temperature of *C. crescentus* (30°C and 34°C) (Fig. 9F). However, there is minimal differences for the Δ*hmrC* mutant: a similar amount of biofilm is formed regardless of the tested temperatures (Fig. 9F). These results suggest that, like the metal response, HmrC plays a role in regulating adhesion under temperatures deviating from the optimal growth temperature. Taken together, we propose that HmrC regulates adhesion in response to environmental stresses, such as the presence of excess copper and extreme temperatures.

## DISCUSSION

From the above results, we postulate that the *hmrABC* gene cluster, along with *hmrX*, encode different elements of a phosphorelay involved in an HfiA-independent regulation of sessile vs. motile lifestyles. We propose that this phosphorelay consists of a transmembrane HHK HmrA, two HPTs (HmrB and HmrX) and a membrane protein (HmrC) at the top of the signaling hierarchy, sensing environmental stress (Fig. 10). HmrA (CC3219 / CCNA_03326) was first reported in a genome-wide study to identify HHKs in *C. crescentus* (51), which reported that a Δ*CC3219* mutant is impaired in swimming through semisolid agar. We show here that, in addition of impacting group swimming, HmrA is also important for adhesion and biofilm formation, by regulating the relative number of cells that produce a holdfast in the population.

**Figure 10:**
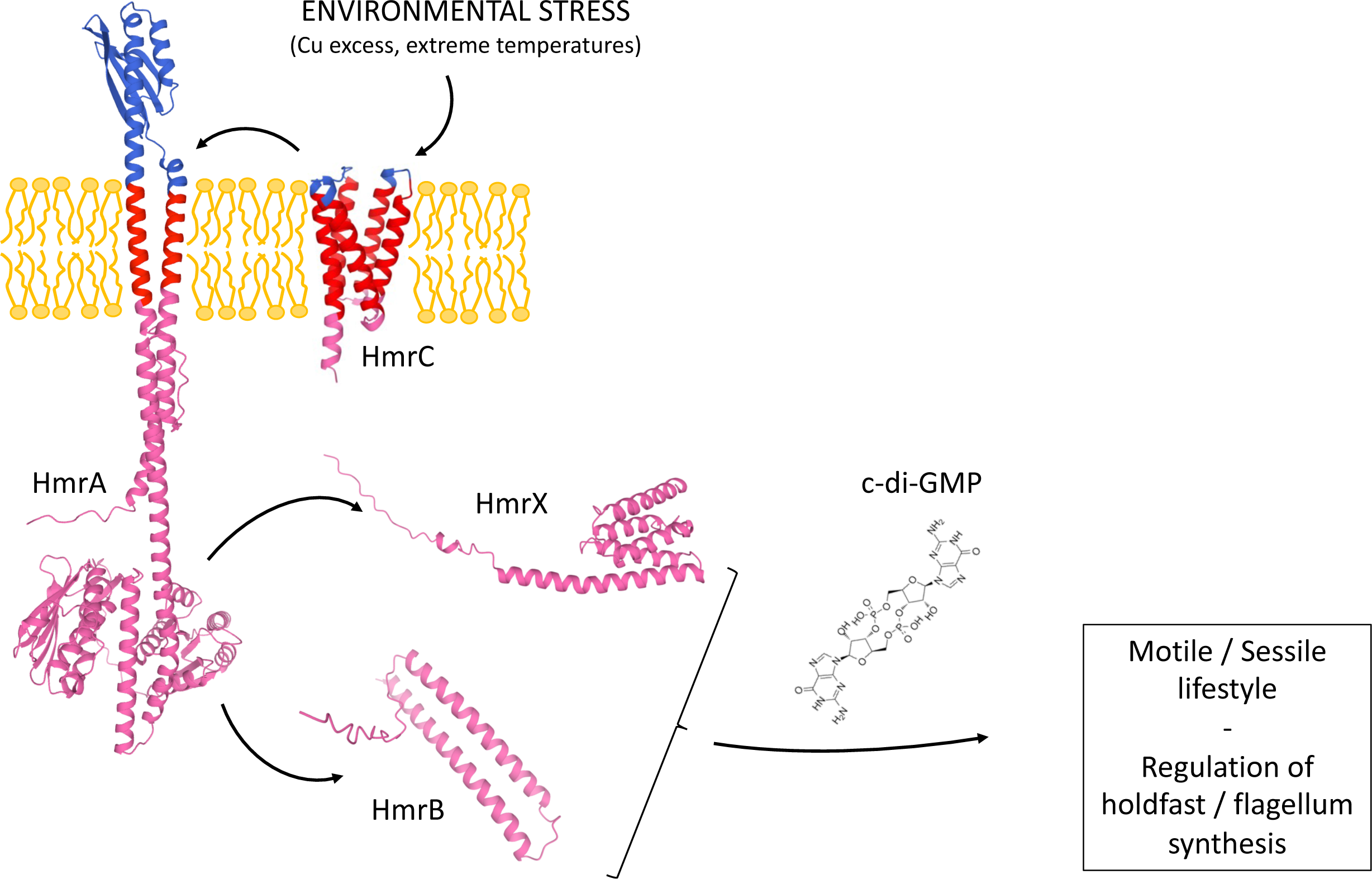
proposed model for the Hmr pathway. Presence of an environmental stress (such as excess copper or extreme temperature) is detected by membrane protein HmrC, which transfers the signal to the HHK HmrA. This signal is transduced to the HTP proteins HmrB and HmrX, which results in a change in c-di-GMP levels via the action of DgcB and PdeA. Changes in c-di-GMP levels modify the regulation of flagellum and holdfast synthesis and dictate the switch from motile to sessile state for each individual cell.

Our work suggests that this system is involved in the balance between the motile and sessile lifestyles. Future biochemical analysis will be required to confirm that the actors of this putative phosphorelay can indeed directly phosphorylate each other, and in which sequence. We showed that residues H286 and D578 in HmrA are crucial for both inhibiting biofilm formation and promoting swimming dispersal. We could not evaluate whether the loss of activity resulted specifically from amino acid substitutions that eliminated catalytic activity, or from an overall structural destabilization. However, since these mutations have been analyzed multiple times in HK research and typically do not destabilize HK structure, we interpret these phenotypes as good indications that *hmrA* encodes a functional HHK. In addition, the most downstream element of the phosphorelay is currently unknown and its identification will be required to complete our knowledge of this pathway.

It is worth noting that the initial screen which enabled the identification of the Hmr pathway revealed two HHKs: HmrA and CCNA_03265 (CC3162) (Fig. 1 and Table 1). Interestingly, CCNA_03265 was also shown to impact group motility through semisolid agar (51). Sequence alignment and structure analysis indicates that the CCNA_03265 HK domain lacks the conserved histidine while the RR domain has retained the aspartate residue (Fig. S1). One possibility would be that the catalytical histidine does not perfectly align with the sequences of other HHKs but is still present in the vicinity. Located a few aa downstream on the same alpha helix, H151 is a reasonable candidate to fulfill this function (Fig. S1). Another possibility would be that CCNA_03265 has no catalytical histidine and that its RR aspartate is phosphorylated by a HPT, such as HmrB or HmrX, or forms a heterodimer with another HHK, such as HmrA. The model of the canonical isolated, linear phosphorelay has prevailed for a long time, but more recent work has shown that phosphorelays can be more complex. For example, in *C. crescentus*, several kinases can phosphorylate the single domain response regulator MrrA (52), which in turn can phosphorylate both the PhyK / PhyR and the LovK / LovR systems known to be involved general stress response (53), cell cycle regulation via HfiA (15), and holdfast production (14). In *P. aeruginosa*, the GacS-GacA TCS (54) interacts with three other HHKs, RetS, LadS, and HHK PA1611, to control the transition between motile and sessile lifestyles depending of the environment (55–57). These examples of regulation cascades involving multiple inputs illustrate the complex molecular mechanisms underlying how bacteria adapt to the different environmental conditions they encounter and how they regulate their lifestyle in response to these changes. Our future work could help to decipher if CCNA_03265 i) is in the same phosphorelay as HmrA; ii) can interact with HmrA; and iii) regulates the transition between motile to sessile lifestyles upon different signals.

### The Hmr pathway regulates motile-to-sessile lifestyle transition independently of HfiA

Several TCS have been shown to regulate holdfast production in *C. crescentus*. For example, the general stress-response PhyR / NepR (58), the LOV (light, oxygen, voltage) blue light photoreceptor proteins LovK / LovR (53), and the RegB / RegA homologs SpdS / SpdR (32) all control holdfast synthesis. They act by regulating HfiA, the major regulator of holdfast synthesis, itself regulated via a multilayered network (15, 19). Interestingly, our work shows that the Hmr pathway operates independently of HfiA for regulation of holdfast production, revealing a novel facet of the highly complex network dedicated to holdfast regulation. A previous ChiP-Seq analysis reported that the cell cycle regulator GcrA, which was shown to bind to the *hfiA* promoter (15), can also interact with the *hmrB* promoter (59). Thus, the connection between GcrA and the Hmr pathway appears as another interesting future direction to investigate to dissect how this cell cycle regulator impacts holdfast production via both the Hmr pathway and HfiA.

### c-di-GMP, produced by DgcB, is involved in the Hmr pathway

We show in this work that the Hmr regulatory pathway might impact the levels of intracellular c-di-GMP, via the diguanylate cyclase DgcB to control both holdfast and flagellum production. Indeed, two different suppressor screens of Δ*hmrA* for gain of swimming through semisolid agar (Fig. 5) identified multiple independent transposon and point mutations in *dgcB*, suggesting that the Hmr pathway and levels of c-di-GMP are linked. This model is in line with the well-established role of c-di-GMP as an important player in the motile to sessile lifestyle transition in *C. crescentus* (60). In addition, our transposon screen for hyper-adhesive mutants (Fig. 1B) enriched for five different mutants in the phosphodiesterase encoding gene *pdeA* (Fig. 1C and Table 1). PdeA affects motility and biofilm formation through c-di-GMP and the activity of DgcB is counteracted by PdeA (13). PdeA is degraded during the motile swarmer to sessile stalked cell transition (61) and DgcB is then able to contribute to increasing c-di-GMP levels in transitioning cells. Stimulation of DgcB activity is achieved by several players at the motile to sessile turning point. First, when swarmer cells encounter a surface in complex medium, pili and the flagellum motor are used as mechanosensors to trigger DgcB to synthesize c-di-GMP, which promotes holdfast synthesis (29, 43). In addition, the <u>C</u>heY-like c-di-GMP <u>e</u>ffectors proteins CleA and CleD are activated by c-di-GMP and interact with the flagellar motor to promote surface contact-stimulated production of holdfast (38). Finally, the flagellar stator MotB and the flagellar signaling suppressor FssA and FssB proteins also trigger DgcB-related c-di-GMP production via the mechanical pathway of holdfast synthesis, regardless of the presence of a surface, via modulation of HfiA expression (42). In those cases, the production of c-di-GMP via DgcB is believed to lead to the activation of the holdfast synthesis protein HfsJ, which binds to c-di-GMP. Our work on the Hmr pathway provides insights into a different regulation mechanism because: i) we performed our experiments in M2X minimal medium where surface contact stimulation of holdfast synthesis is not active (30); and ii) we showed that HfiA is not involved in the Hmr pathway. Future work could provide more insight into whether HfsJ is the final player in the Hmr regulation cascade.

### The Hmr pathway is important for tuning biofilm formation in response to environmental stresses

Our results suggest that the Hmr pathway is important for adapting the sessile-to-motile transition in response to environmental cues, as HmrC is involved in the response to environmental stresses such as presence of excess metals or non-optimal temperatures (Fig. 9E-F). *hmrC* was shown to be upregulated in presence of metals and is part of the UzcR regulon, which is involved in U, Zn and Cu sensing (49). We tested various metals, namely Cu, Zn, Co, Fe, Mn, and Ni. In the tested conditions, only Cu excess produced a significant alteration of the biofilm phenotype in WT cells. Cu stress in *C. crescentus* depends on both the growth medium and the cell type. Indeed, it was shown that cells are overall more sensitive to excess Cu when grown in defined medium than in rich medium PYE (62). A recent RNAseq study showed that *C. crescentus* reacts to Cu excess in M2G by overexpressing machineries dealing with Cu detoxification (PcoAB), oxidative stress, protein misfolding/rearrangement and chelation by cysteine and arginine. In the same study, it stands out that *hmrC* is the only gene of the *hmr* set that is consistently upregulated by addition of Cu (2.4x in stalked cells; 4.1x in swarmer cells). Other *hmr* genes displayed either no change or a 2-fold downregulation upon Cu addition. Considering our results, HmrC is an important factor in managing the response to excess copper. We show here that biofilm formation in *C. crescentus* WT is decreased by the addition of Cu, while it remains steady in a Δ*hmrC* mutant, suggesting that this protein is important to modulate biofilm adhesion under Cu stress. Apart from the growth medium, the cell type of *C. crescentus* determines the response to Cu excess. While swarmer cells preferentially escape excess copper by negative chemotaxis, stalked cells are seemingly more impacted by Cu stress and rely on detoxification mechanisms (primarily PcoAB) (63). Such dimorphism was postulated to condition a fast bimodal response to Cu stress. Since HmrC, and the Hmr pathway in general, appears to determine the proportion of motile vs. sessile cells in a mixed population, we hypothesize that the Hmr pathway, with HmrC reponding to Cu levels, is a molecular mechanism underlying this bimodal response by buffering dimorphism.

*C. crescentus* has an optimal growth temperature of 30°C. Our results show that it produces approximately 5 times more biofilm at 15°C and 37°C and that this difference depends on HmrC (Fig. 9F). Interestingly, *hmrC* is upregulated during cold shock and cold shock affects metal and ion homeostasis (50). It is not clear at this stage whether HmrC is involved in sensing broad environmental stress, including metal-induced *and* cold-induced stress, or whether it is involved in sensing environmental metals and its role in cold sensing is a consequence of the disturbance of metal homeostasis. Temperature changes can destabilize the bacterial membrane by changing its fluidity. In the case of cold shock, the main signal that triggers adaptation is the sensing of membrane rigidity by transmembrane histidine kinases, as described in a broad range of organism such *Escherichia coli*, *Bacillus subtilis*, *Salmonella typhimurium*, *Yersinia pestis* and *Synechocystis* PCC6803 (64). Based on our results, we hypothesize that the membrane protein HmrC is required for sensing temperature change, which conditions the proportion of biofilm forming cells as a function of the temperature. Interestingly, deletion of *hmrC* comes at a high fitness cost according to a genome wide gene essentiality study in *C. crescentus* (65). Overall, this could point to HmrC as a crucial element for sensing environmental stresses. We can then speculate that this protein sends a signal to HmrB and / or HmrX, via HmrA to eventually modulate the transition from the motile to the sessile lifestyle, in a c-di-GMP dependent manner, and ensures that cells settle in favorable environments.

## MATERIAL AND METHODS

### Bacterial strains and growth conditions

All bacterial strains used in this study are listed in Table S1. *Escherichia coli* strains used for cloning were grown in LB medium at 37°C, in the presence, when necessary, of antibiotics at the following final concentrations: kanamycin 50 µg ml^-1^ (for pMR10 constructs), gentamycin 20 µg ml^-1^ (for pTNS3 constructs), and tetracycline 15 µg ml^-1^ (for pRKlac290 constructs). *C. crescentus* strains were grown at 30°C using M2 minimal medium with 0.2% xylose (M2X) (66) in liquid, and using Peptone Yeast Extract (PYE) (67) + 15 g l^-1^ bactoagar (Difco) plates. When necessary, antibiotics were added to *C. crescentus* cultures at the following final concentrations: kanamycin 20 µg ml^-1^ (for pMR10 constructs), gentamycin 50 µg ml^-1^ (for pTNS3 constructs), and tetracycline 2 µg ml^-1^ (for pRKlac290 constructs).

### Plasmid construction and cloning procedures

All plasmids were cloned using standard molecular biology techniques. PCR was performed using *C. crescentus* NA1000 *hfsA*^+^ WT genomic DNA as template. Sequences of the primers used are available upon request.

In-frame deletion mutants were obtained by double homologous recombination, as previously described (68). Briefly, 500-bp fragments from the upstream and downstream regions of the gene to be deleted were amplified by PCR. PCR fragments were gel-purified and then digested by *Hind*III and *BamH*I or *BamH*I and *EcoR*I for upstream or downstream fragments, respectively. Purified digested fragments were then cloned into the suicide vector pNPTS139 that had been digested by *EcoR*I and *Hind*III. The pNPTS139-based constructs were transformed into *E. coli* DH5a cells and then introduced into *C. crescentus* by electroporation. The two-step recombination was carried out using sucrose resistance and kanamycin sensitivity (68). Then, the mutants were verified by sequencing to confirm the presence of the deletion.

The pMR10-*hmrA* and pMR10-*hmrB* constructs were created as follows. The genes and their putative native promoter (approximately 500 bp upstream of the start codon) were amplified by PCR, gel-purified and then digested by *EcoR*I and *BamH*I. Purified digested fragments were ligated into pMR10 that had been digested with *EcoR*I and *BamH*I. Ligation products were used to transform *in E. coli* DH5α cells by heat shock. Once plasmids were isolated from *E. coli* and the sequence of the insert was confirmed through sequencing, plasmids were electroporated into *C. crescentus. C. crescentus* cells harboring the pMR10 derivatives were screened for kanamycin resistance. The point-mutation derivatives of these strains were generated using oligos containing the appropriate point mutation.

The complementation of mutants in *hmrA, hmrB, hmrC*, and *hmrX* was performed by inserting each gene with its promoter at the Tn7 *att* site. Briefly, a region consisting of the gene of interest was amplified with flanking regions to preserve the promoter and terminator sequences (approximately 300bp upstream and 100-200bp downstream). The PCR product was subsequently inserted into pUC18-mini-Tn7-LAC (69). The resulting plasmid was introduced in the *C. crescentus* strain of interest along with the helper plasmid pTNS3 (70) to insert the gene of interest under its native promoter at the Tn7 *att* site in the chromosome of *C. crescentus*. This insertion contains a cassette that confers resistance to gentamycin, which enables selection.

*C. crescentus* NA1000 *hfsA^+^* derivatives harboring the stable miniTn7*dsred* were constructed by ΦCR30 mediated transduction (71) from CB15::miniTn7*dsred* (YB4788) lysate as previously (72). The NA1000 *hfsA^+^ divJ-cfp / pleC-yfp* strains were constructed by ΦCR30 mediated transduction (71) from CB15N *divJ-cfp / pleC-yfp* (73) lysate.

### Motility assays through semisolid media

Motility assays were performed using semisolid agar plates. Plates were poured using M2 medium supplemented with 0.2% (w/v) xylose and 0.4% (w/v) noble agar (Difco) and allowed to sit overnight at room temperature. Cells from colonies grown on regular PYE agar were stabbed into the soft agar and incubated in a humid chamber at 30°C for 3 days or room temperature for 5 days. The diameter of the swimming ring formed by each tested strain was measured manually and normalized to that of WT (with empty vector when appropriate).

### Biofilm assays

Biofilm assays in 24-multiwell plates were performed as described previously (72). Bacteria were grown to mid-log phase (OD_600_ of 0.3 - 0.6) in M2X and diluted to an OD_600_ of 0.05 in the same medium. 500 µl of freshly diluted cells were placed in each well. Plates were incubated at 30°C for 24 h or otherwise as mentioned in the text. Biofilms attached to the inside surface of the wells were quantified as follows: wells were rinsed with distilled H_2_O to remove non-attached bacteria, stained using 0.1% crystal violet (CV), and rinsed again with dH_2_O. The CV from the stained attached biomass was eluted using 10% (v/v) acetic acid and was quantified by measuring absorbance at 600 nm (A_600_). Biofilm formation was normalized to A_600_ / OD_600_.

### Holdfast quantification using fluorescently labeled WGA lectin

The number of cells harboring a holdfast in mixed populations was quantified by fluorescence microscopy. Holdfasts were detected with AlexaFluor 488 conjugated wheat germ agglutinin (AF488-WGA) since WGA binds specifically to the N-acetylglucosamine residues present in the holdfast (74). Early exponential-phase cultures (OD_600_ of 0.2 - 0.4) were mixed with AF488-WGA (0.5 µg/ml final concentration in water). One microliter of WGA-stained cells was spotted on a 1.5-mm glass coverslip and covered with an agarose pad (1% SeaKem LE Agarose dissolved in dH_2_O). Holdfasts were imaged by epifluorescence microscopy. The number of individual cells with a holdfast was calculated manually from microscopy images.

### *hfiA* expression assay using β-galactosidase reporter

Strains bearing a transcriptional reporter plasmid for the *hfiA* gene promoter fused to *lacZ* (15) were inoculated into 5 ml of M2X containing 2 μg/ml tetracycline and incubated at 30°C overnight. Cultures were then diluted in the same culture medium to an OD_600_ of 0.05 and incubated until an OD_600_ of 0.15 – 0.3 was reached. β- galactosidase activity was measured colorimetrically as described previously (75). 200 µl of cells were added to 600 µl of Z buffer (60 mM Na_2_HPO_4_, 40 mM NaH_2_PO_4_, 10 mM KCl, 1 mM MgSO_4_, 50 mM ß-mercaptoethanol), 50 µl of chloroform, and 25 µl of 0.1% SDS. Two-hundred microliters of substrate *o*-nitrophenyl- β-D-galactoside (ONPG, 4 mg/ml) was then added and time was recorded until development of a yellow color. The reaction was stopped by adding 400 µl of 1M Na_2_CO_3_. Absorbance at 420 nm (A_420_) was measured and the Miller Units of β-galactosidase activity were calculated as (A_420_ x 1000)/((OD_600_ x *t*) x *v*), where *t* is the incubation time in minutes after addition of ONPG, and *v* is the volume of culture (in ml) used in the assay. The β-galactosidase activity of WT plac290 (empty vector control) was used as a blank sample reference.

### Holdfast synthesis timing by time-lapse microscopy

Cell division and holdfast synthesis timing were observed in live cells on agarose pads by time-lapse microscopy as described previously (30). A 1 µl aliquot of exponential-phase cells (OD_600_ of 0.4–0.7) was placed on top of a pad containing 1% SeaKem LE Agarose diluted in M2X and AF488-WGA (0.5 µg/ml final) sealed under a coverslip with valap. Time-lapse microscopy images were taken every 2 min for 8 h. Time-lapse movies were visualized in ImageJ (76) to manually assess the time of cell division, as *t* = 0 when a cell newly divides, the time of holdfast production, as the first frame where the fluorescent lectin signal is detected, and the time of new division.

### Imaging and labeling of flagellum filaments

Strains used to visualize flagella by fluorescence microscopy harbored a cysteine-knock in the major flagellin *fljK* mutants were used for flagellum filament staining (30), as well as a miniTn7*dsred* insertion, for cell body visualization (72). Cell cultures were grown to mid-exponential phase (OD_600_ = 0.3–0.4) in M2X medium. 100 µl of culture was mixed with 10 µl of AF488 conjugated maleimide (AF488-mal, 50 µg/ml in water) for a final dye concentration of 5 µg/ml and incubated for 5 min at room temperature. Cells were centrifuged at 5,000 *g* for 1 min, washed with 1 ml of sterile water to remove excess dye and shed flagella and resuspended in 50 µl of sterile water. One microliter of the labeled culture was added to a coverslip and imaged under a 1% SeaKem LE Agarose diluted in water. Cell bodies and flagella were imaged using red and green fluorescence microscopy settings. The number of cells bearing a flagellum in the mixed population was quantified for microscopy images using ImageJ (76).

### Epifluorescence microscopy

Epifluorescence microscopy was performed using an inverted Nikon Ti2 microscope with a Plan Apo 60X objective, a GFP/DsRed filter cube, an Andor iXon3 DU885 EM CCD camera, and Nikon NIS Elements imaging software. Image analysis was performed using in the Nikon NIS Elements or Image J (76) analysis software built-in tools.

### Nanoscale Atomic Force Microscopy (AFM) imaging

100 µl of cell cultures grown to mid-exponential phase (OD_600_ = 0.3–0.4) in M2X medium were diluted in 900 µl sterile water. 10 µl were spotted to a clean glass coverslip and allowed to dry at room temperature. AFM imaging was caried out in air at room temperature, using a Bioscope resolve AFM (Bruker) using the peak-force tapping mode, using silicon cantilevers with a nominal spring constant of around 0.4 N/m and a nominal tip radius of 2 nm (ScanAsyst-Air, Bruker).

### Cell synchronization, initial attachment of *C. crescentus* single cells to surfaces, and motile cell swimming speed quantification

Small scale synchronization of *C. crescentus* cells was performed as described previously (77) with some modifications. Overnight cultures in M2X were diluted 10 times in 30 ml fresh M2X, and incubated shaking at 50 rpm at room temperature in a 150 mm diameter Petri dish. After overnight incubation, the medium was removed and the monolayer biofilm formed in the dish was thoroughly washed with sterile water. 30 ml of fresh M2X medium was added to the dish and the incubation was carried out for 4 h under the same conditions. The dish was then rinsed 10 times using 10 ml of fresh M2X medium to remove unattached cells from the biofilm. 1 ml of M2X medium was added to the dish and the incubation was carried out 5 min under the same conditions to release the newly born swarmer cells. The homogeneity of the synchronized swarmer population was verified by microscopy.

Initial attachment of *C. crescentus* cells to surfaces was recorded by dark-field microscopy as described previously (72). One microliter of synchronized swarmer cells was observed by dark-field microscopy (10X objective). A stack of 100 frames (100 ms exposure time) was recorded using RAM capture every minute for one hour. The maximum intensity of the 20 first frames of each stack was calculated to visualize motile cells with a swimming trajectory, using the built-in functions in ImageJ. The number of swimming trajectories in the field of view was quantified manually and the value calculated for each sample at t=0 was set as 100%.

RAM captures stacks recorded were also used to determine trajectories and velocities of swimming cells. The entire tracks (1 second movies) were analyzed using the MicrobeJ plugin (78) in ImageJ. For each strain, more than 500 cell tracks were analyzed.

### Chemotaxis assays

Chemotaxis assays were performed as previously described (39). Plates were poured using M2 (no carbon added) + 0.2% noble agar. A sterile 500 µm diameter filter paper disk containing 50 µl xylose (20% solution) was placed in the center of the place, and cells were spotted at 1 cm away from the chemoattractant-filled disk. Plates were incubated at 30°C for 48h. Chemotaxis rings were imaged using a ChemiDoc imaging system (Biorad), and the relative pixel intensity profile through the center of the chemotaxis ring was calculated for each strain.

### *Mariner* transposon mutagenesis

For *Mariner* transposon mutagenesis, the pFD1plasmid bearing the *Himar1*-based minitransposon (22) was mated into NA1000 *hfsA*+ WT or Δ*hmrA*. Mutants bearing the transposon were selected on PYE + kanamycin (20 µg/ml) and screened for hyper-adhesion or improved motility through semisolid agar as described below.

Genomic DNA from mutants of interest was isolated and subsequently used as a template for inverse PCR to identify the insertion site of the *Mariner* transposon. 1μg of genomic DNA was digested with 5 units of Sau3AI. The digested DNA was then self-ligated with T4 DNA ligase. 25-150 ng of the self-ligated genomic DNA was mixed with the primers (ACGGTATCGATAAGCTTGATATCGA and CAGAGTTGTTTCTGAAACATGGCA) and PCR amplified. PCR fragments were then gel purified and sequenced using amplification from the transposon primer at the Institute for Molecular and Cellular Biology, Indiana University, Bloomington, USA.

### Screen for hyper adhesive mutants

Wild-type *C. crescentus* NA1000 *hfsA^+^* cells were randomly mutagenized with the *Mariner* transposon as described above. Over 15,000 colonies were selected on PYE + Kanamycin plates, providing greater than 95% genome coverage (79, 80). Colonies were pooled in sets of approximately 150 in 3 ml M2X medium and incubated overnight in plastic 12-multiwell plates with a 18 x 18 mm glass coverslip added vertically in each well. After incubation, the cells that were bound to the coverslips were used as inoculation for a new culture: coverslips were washed carefully with fresh M2X medium to remove any loosely attached cells, added to a new well containing 3 ml of fresh M2X and incubated for 3 h. After incubation, new sterile glass coverslips were added to the new culture in the wells, and the process was repeated. After two enrichments the cultures were streaked for single colonies. Approximately 300 colonies were randomly picked and subjected to a secondary screen to identify mutants with more than 200% biofilm formation compared to the WT control.

### Screen for Δ*hmrA* suppressor mutations on semisolid agar plates

Two different types of approaches were taken to isolate suppressor mutations that would restore the motility defect of the Δ*hmrA* mutant in semisolid agar.

i) The NA1000 *hfsA+* Δ*hmrA* mutant was mutagenized with *Mariner* transposons as described above. Approximately 26,000 colonies were selected on PYE + Kanamycin plates. This sample size provided greater than 97% genome coverage (79, 80). The colonies were pooled together into a set of 15 pools and normalized to OD600nm = 0.3. Two microliters from each of the normalized pools was stabbed into a PYE plate containing 0.3% agar, and incubated at 30°C. After 48 hours, cells from edge of spreading flares were streaked to obtain single colonies. Only one single flare was selected from each initial semisolid plate, to avoid isolating siblings, yielding 15 independent suppressors. A fresh inoculation of each isolated suppressor strain in semisolid agar confirmed their phenotype (swimming ring greater than Δ*hrmA* sized rings). To validate that the phenotype suppression was due to the *Mariner* transposon insertion, ΦCR30 lysates of the 15 mutants were created for transduction as described below. These lysates were used to transduce the *Mariner* transposon insertion into the Δ*hmrA* mutant background. An isolated colony from each of the 15 transductants was inoculated into semisolid agar plates, 14 out of the 15 transduced mutants retained the increased swimming diameter and were studied further. Genomic DNA was extracted and inverse PCR was performed to identify the transposon insertion site in each mutant.
ii) Random spontaneous motile suppressors were isolated as follow. Eight separate cultures of *NA1000 hfsA+* Δ*hmrA* were grown in 5 ml PYE overnight and streaked on PYE plates. One isolated colony from each plate was stabbed into a semisolid PYE plate and incubated at room temperature. After 96 hours, cells were isolated from the edge of different flares and streaked on fresh PYE plates. The swimming phenotype of the isolated motility suppressors was confirmed on semisolid PYE plates and only the strains exhibiting a swimming ring greater than that of Δ*hmrA* were carried further for identification. For each suppressor mutant, the *dgcB* gene was first sequenced, and only strains with a WT copy of *dgcB* were sent for Illumina sequencing (3 out of 11 mutants).

### DNA transduction by ΦCR30 phage lysate

Phage lysates were prepared as follows (71). Donor strains of *C. crescentus* were grown overnight at 30°C in PYE. 100 μl of a 10^-3^ dilution of live ΦCR30 phage and 200 μl of the overnight donor culture were added to the molten PYE top agar (0.5% agar), mixed and poured onto the surface of a pre-heated PYE plate. After overnight incubation at 30°C, 5 ml of fresh PYE was added on to the surface of the plate and incubated overnight at 4°C. The remaining PYE liquid was removed from the plate and transferred to an empty 100 mm Petri dish and irradiated under a germicidal UV lamp for 5 minutes. After irradiation, 200 μl chloroform was added to samples transferred to a glass screw cap tube and stored at 4^°^C protected from light.

Transductions were done by mixing 500 µl of overnight *C. crescentus* recipient culture, 50 μl of UV-irradiated phage lysate (prepared as described above), and 475 μl of fresh PYE. The mixture was incubated for 4 h at 30°C and plated on a PYE + kanamycin plate, to select for transductants harboring a DNA fragment including the *Mariner* transposon.

### ΦCbK phage sensitivity assay for presence of pili

Overnight cultures to be tested were normalized to OD600 = 0.6 and serially diluted in steps of 100x to a final concentration of 10^-8^. 100 μl of ΦCbK phage solution was uniformly absorbed onto the surface of a PYE plate, and 10 μl of culture dilutions spotted on the plate and allowed to dry prior to incubation at 30°C. The number of colonies were recorded after for two days of incubation.

### Quantitative PCR to evaluate *Cori/ter* ratios

The relative chromosomal presence of the *Cori* and *ter* regions were determined as described previously (81) with a few modifications. Briefly, the following primer set was used: Cori_fwd (5′-CGCGGAACGACCCACAAACT-3′) and Cori_rev (5′-CAGCCGACCGACCAGAGCA-3′) to amplify a region close to the origin (*Cori*); Ter_fwd (5′-CCGTACGCGACAGGGTGAAATAG-3′) and Ter_rev (5′-GACGCGGCGGGCAACAT-3′), to amplify a region close to the terminus (*ter*). Cells were grown in M2X medium and harvested in exponential phase. Genomic DNA was extracted using the Monarch genomic DNA extraction and purification kit (New England Biolabs Inc.). Region of *Cori* and *ter* were quantitatively amplified using the Sybr green-based Luna Universal One-step RT-qPCR kit (New England Biolabs Inc.), without the reverse transcriptase enzyme, in a QuantStudio 3 thermocycler (Applied Biosystems, Thermo Fisher Scientific). All procedures were performed according to the manufacturer recommendations. qPCR reactions were performed in triplicate. Efficiencies of Cori_fwd/Cori_rev and Ter_fwd/Ter_rev primer pairs were evaluated from a 5-fold dilution series of 8 different concentrations. Relative abundance of *Cori* to *ter* was calculated using the Pfaffl method, which considers the respective primer set efficiencies (82). Cori/ter ratio of Δ*hmrA* was normalized to that of WT. Three independent biological replicates were used for this quantification.

### Cell-cycle specific protein localization by fluorescence microscopy

To assess cell cycle timing, cells expressing fluorescent fusions to ParB, DivJ and PleC proteins were tracked by microscopy. The localization and duplication of the chromosome partitioning protein ParB was tracked by time lapse microscopy using strains harboring a *parB::gfp* fusion (83). To ensure that both WT and Δ*hmrA* strains were subject to the same conditions during the experiment, one strain was labelled with dsRed and the strain that was labelled was switched for each independent replicate. One µl of a mixture of dsRed-labelled/unlabelled cells (OD_600_ = 0.4 – 0.6) was placed on top of a pad containing 1% SeaKem LE Agarose diluted in M2X and sealed under a coverslip with valap. Time-lapse microscopy images were taken every 2 min for 8 h. Time-lapse movies were visualized in ImageJ (76) to manually assess the timing of ParB duplication (swarmer to stalked cell differentiation), with *t* = 0 defined as the time of division.

The localization of PleC and DivJ was determined in cells harboring both a DivJ-CFP and a PleC-YFP fusion (73), which were synchronized as described above. Synchronized cells were harvested from birth from a synchrony petri dish every 10-15 minutes and imaged under a 1% SeaKem LE Agarose diluted in water. 10 random fields of view were taken for phase contrast and CFP and YFP fluorescence for each time point (three independent replicates). Quantification of the number of cells per field of view with a localized PleC-YFP and DivJ-CFP was performed manually.

### Biofilm formation assay in response to the presence of metals or temperature change

In order to study metal dose-response of *C. crescentus*, strains were grown overnight in M2X at 30°C with shaking at 180 rpm. Starter cultures were diluted to an OD_600_ of 0.05 in 500 µl of M2X containing various amount of metal salts (0-10 µM for CuSO_4_ and 0-100 µM for ZnSO_4_) in 24-well plates. Plates were incubated statically overnight, and biofilm assay was performed as describe above.

In order to study response to temperature, the procedure was the same except that no metal was added to the medium and plates were incubated at various temperatures, as indicated.

## ACKNOWLEDGEMENTS

This work was supported by a Canada 150 Research Chair in Bacterial Cell Biology to Y.V.B. Early stages of this work were supported by grant R35GM122556 to Y.V.B. from the National Institutes of Health. We thank Marylise Duperthuy for the use of the qPCR machine. We are also grateful to all the members of the Brun lab for feedback on this research and for critical review of the manuscript.

---

**Figure S1:**
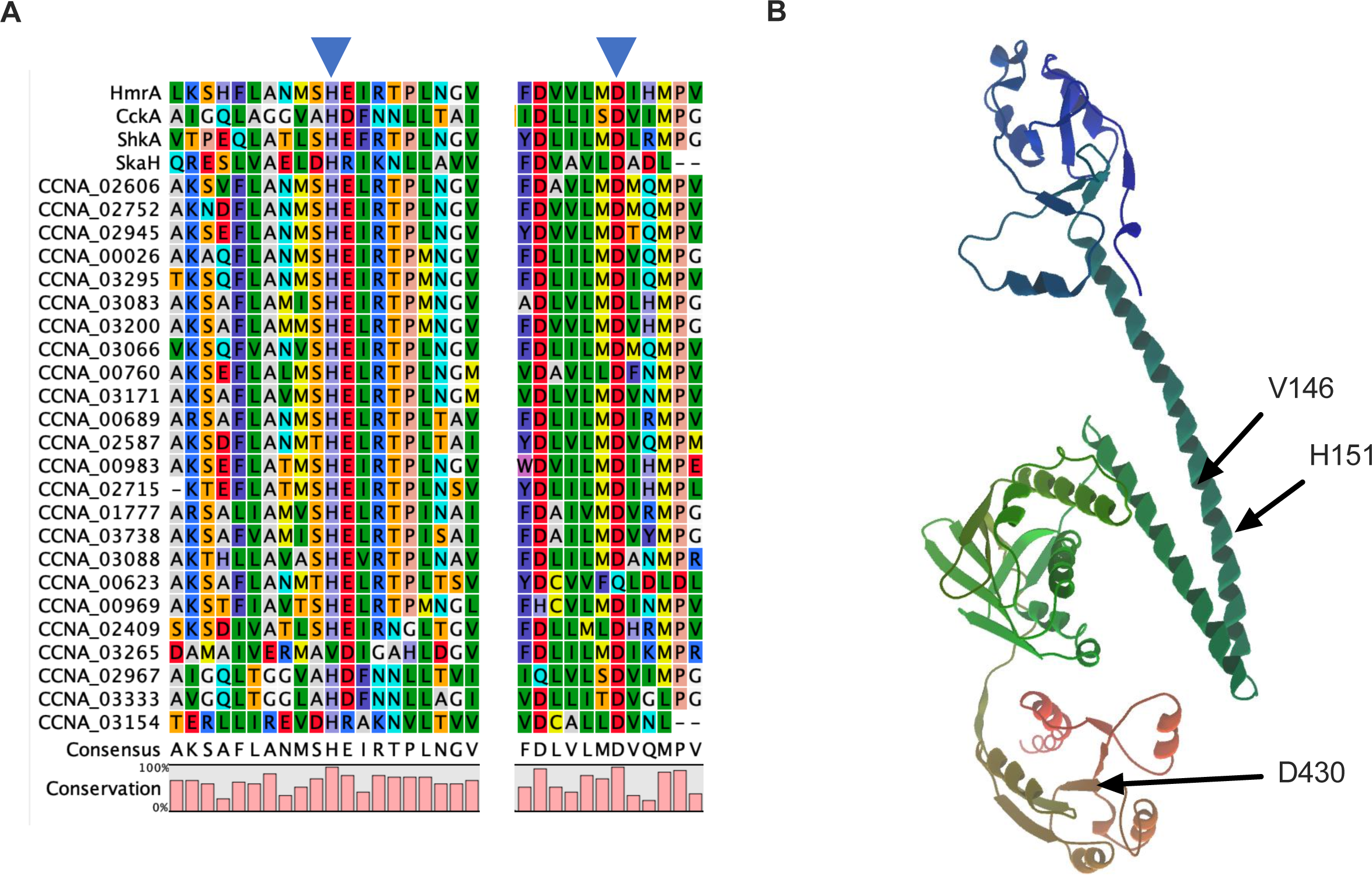
structural analysis of HHKs identified in the *C. crescentus* genome. **(A)** Protein sequence alignment for the active sites of the 28 predicted HHK present in the *C. crescentus* genome, including the experimentally verified CckA, ShkA and SkaH. Conserved putative catalytical residues, corresponding to H286 and D578 in HmrA, are highlighted with arrows. **(B)** Prediction of tridimensional structure of CCNA_03265 encoded HHK, using AlphaFold. V146 is the residue that aligns with the putative catalytic histidine of other HHKs (cf. panel A). H151 is a candidate as catalytic histidine residue. Putative catalytic aspartate, D430, is also displayed.

**Figure S2:**
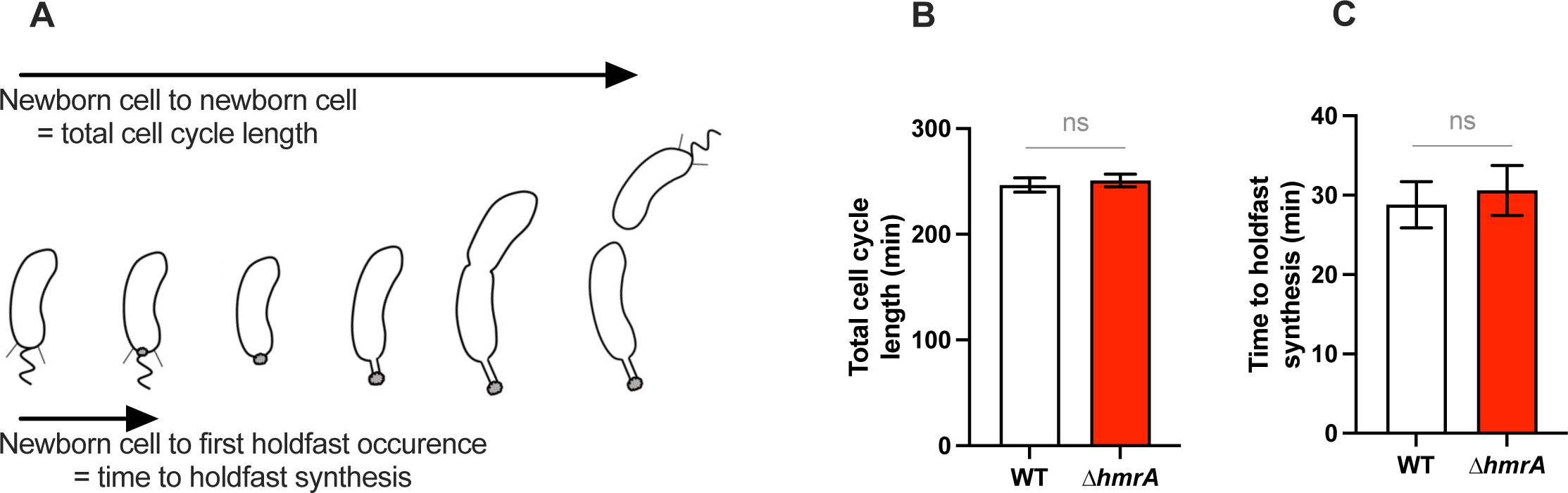
the cell cycle and timing of holdfast synthesis is not affected in a Δ*hmrA* mutant. **(A)** Representation of total cell cycle length and time to holdfast synthesis, which are measured in the panels B and C. **(B)** Timing of total cell cycle length and **(C)** Timing of holdfast synthesis by newly divided swarmer cells on M2X agarose pads containing AF488-WGA. The timing between when a new cell divides and when the holdfast first appears on that same cell (B) and when that cell completes an entire cell cycle (A) are recorded. Around 50 cells were counted in three independent replicates. Statistical comparisons are calculated using unpaired *t*-tests (ns = not significant).

**Figure S3:**
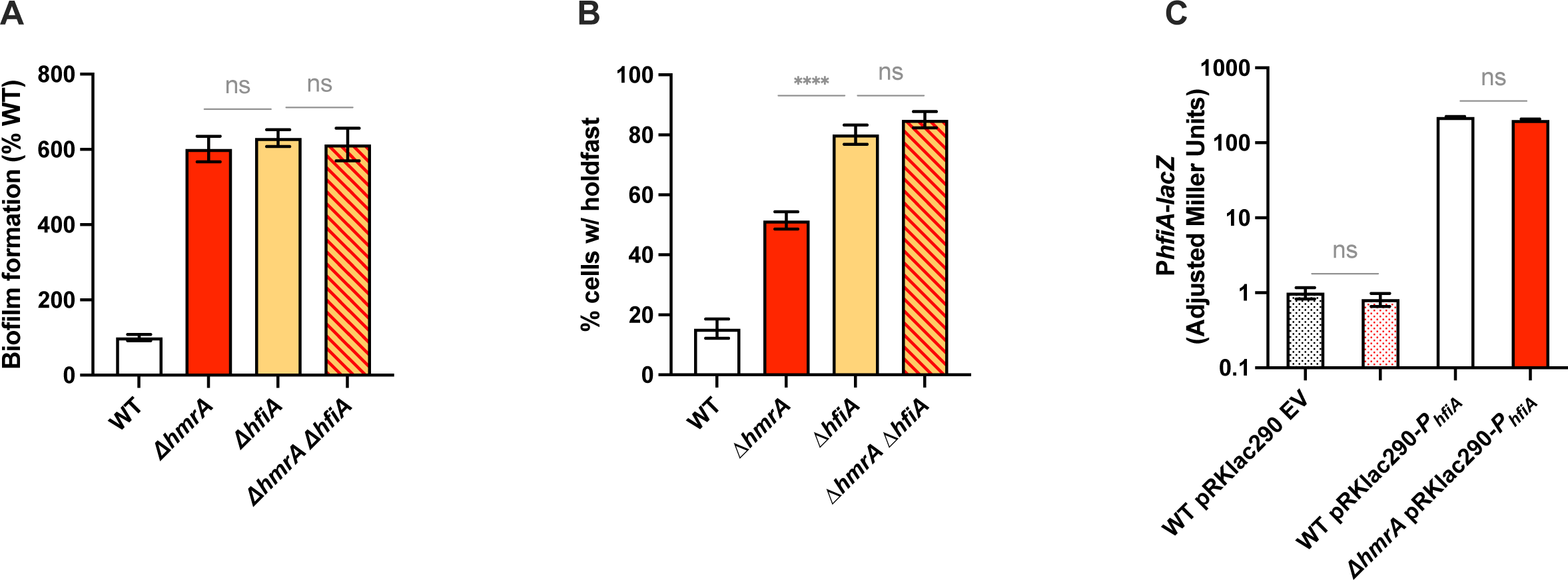
the holdfast inhibitor HfiA is not involved in the HmrA mediated holdfast regulation pathway. **(A)** Biofilm formation. Results are expressed as a percentage of biofilm formed by each strain compared to WT set at 100%. Results are given as the average of three independent experiments, each run in triplicate, and the error bars represent the Standard Error of the Mean (SEM). Statistical comparisons are calculated using unpaired *t*-tests (ns: not significant)**. (B)** Quantification of cells harboring a holdfast in mixed populations. The results represent the average of three independent replicates (more than 250 cells per replicate) and the error bars represent the SEM. Statistical comparisons are calculated using unpaired *t*-tests (ns: not significant; **** *P* < 0.0001)**. (C)** ß-galactosidase activity of the P*_hfiA_*-*lacZ* transcriptional fusions in WT and Δ*hmrA* strains. The results represent the average of three independent cultures (assayed on 3 different days) and the error bars represent the SEM. Statistical comparisons are calculated using unpaired *t*-tests (ns: not significant). For each replicate, the **r**esults are normalized to the baseline ß-galactosidase activity from the WT pRKlac290 EV (empty vector) strain (set to 1).

**Figure S4:**
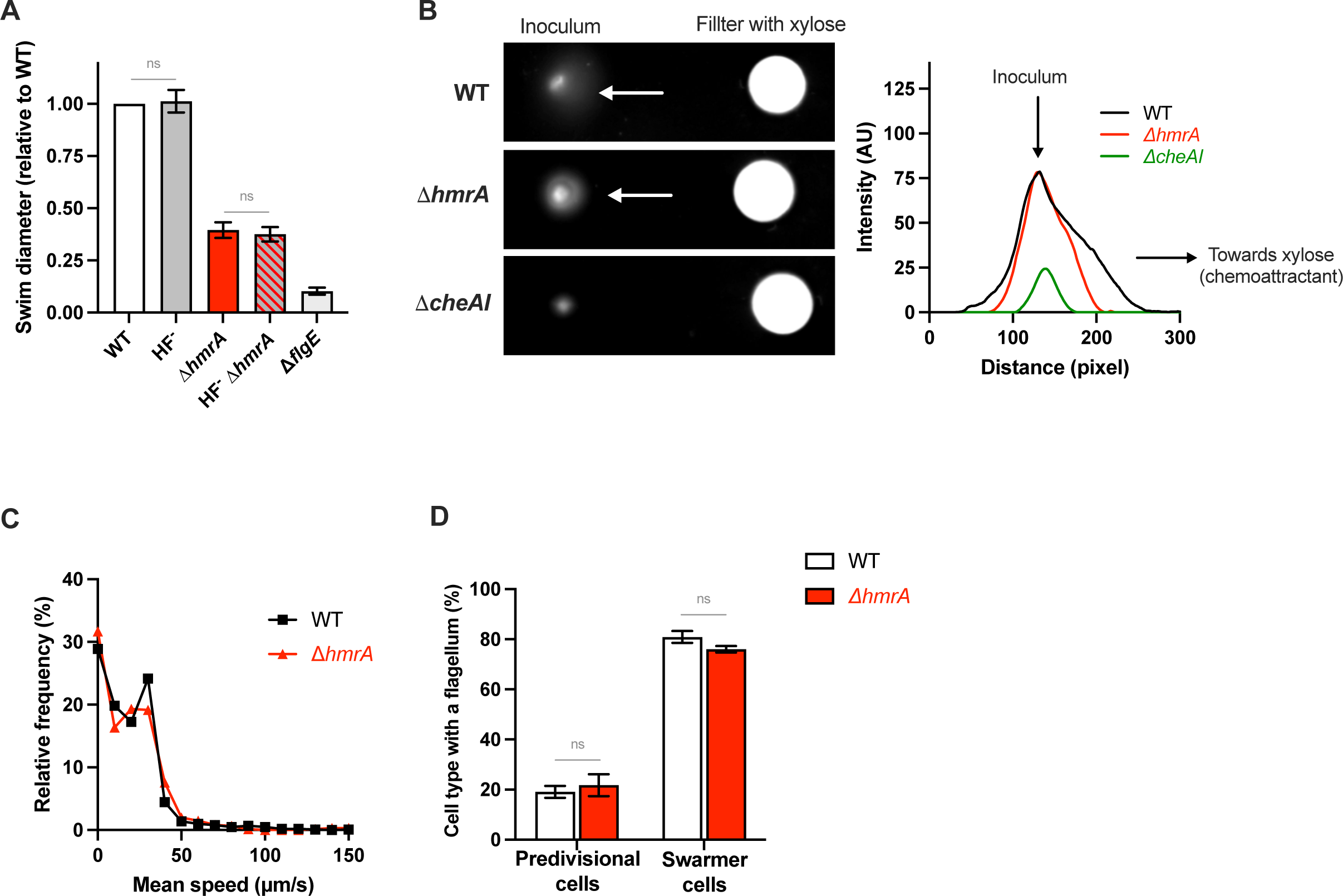
holdfast is not required for the impaired swimming through semisolid medium of the *ΔhmrA* mutant, and the *ΔhmrA* mutant is not impaired for chemotaxis or for motility in liquid. **(A)** Motility assays in semisolid agar. Results are normalized to WT (NA1000 *hfsA*+) ring diameter measured on the same plate (set to 1). The HF^-^ strain (NA1000) does not produce holdfasts. Bar graphs indicate the mean of three independent replicates, and error bars represent SEM. Statistical comparisons are calculated using unpaired *t*-tests (ns: not significant)**. (B)** Chemotaxis assays. Strains are inoculated by spotting into M2 (no carbon) + 0.2% noble agar plate and incubated for 48h. A filter paper containing xylose was placed at a similar distance for the inoculum for each tested strain, to act as a chemoattractant. The relative pixel intensity profile through the center of the chemotaxis ring in a representative image is shown on the right. **(C)** Frequency distribution for swimming run mean speeds. **(D)** Proportion of swarmer and predivisional cells harboring a flagellum. The results are calculated as a percentage of total flagellated cells for each strain (set as 100%) and represent the average of at least three independent replicates (more than 200 cells per replicate). The error bars represent the SEM. Statistical comparisons are calculated using unpaired *t*-tests (ns: not significant).

**Figure S5:**
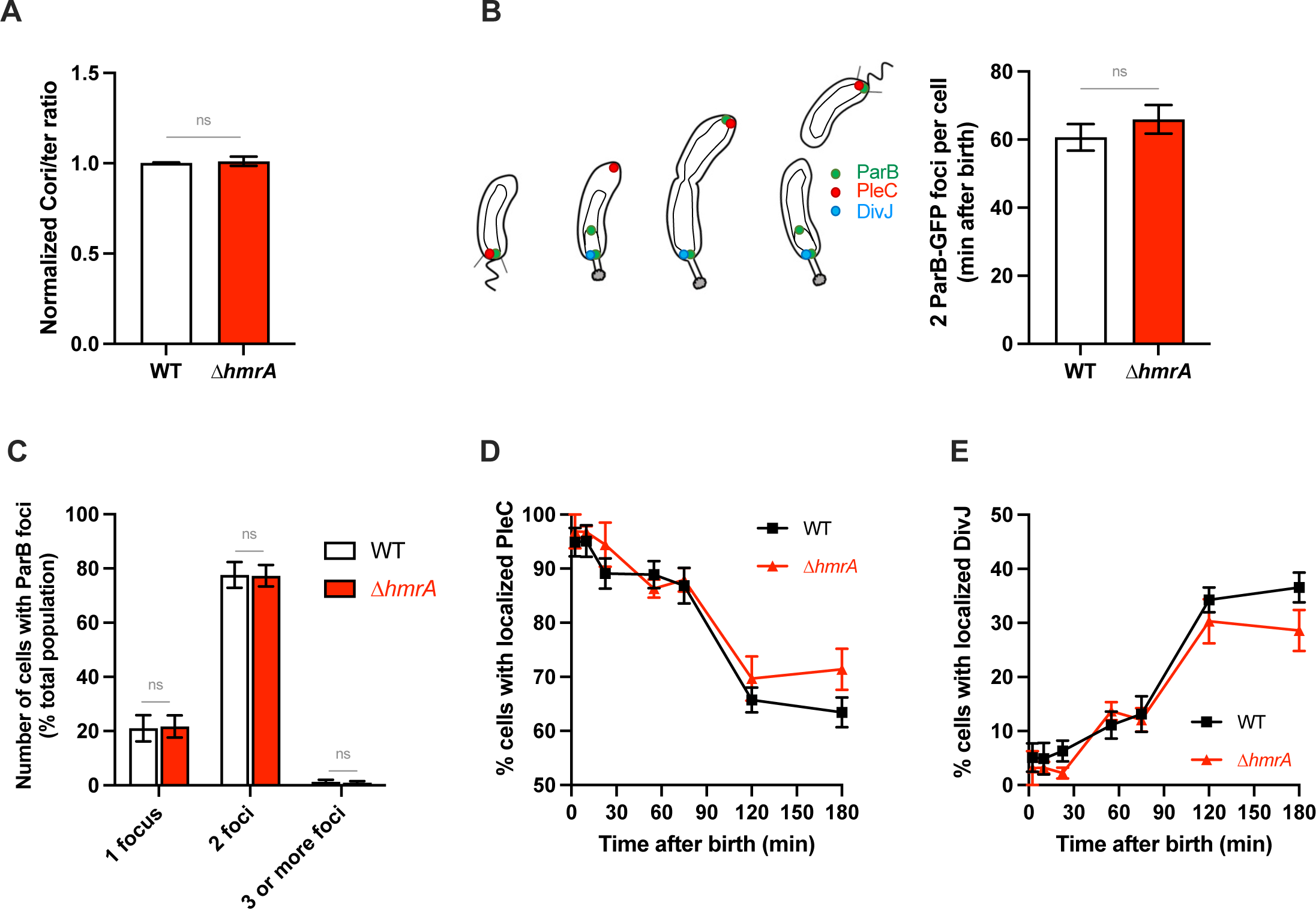
HmrA does not regulate *C. crescentus* cell differentiation and chromosome segregation. **(A)** Relative *Cori/ter* ratio measured by qPCR. Results are given as the average of three independent replicates, normalized for WT in each replicate, and the error represents the SEM. Statistical comparisons are calculated using unpaired *t*-tests (ns: not significant)**. (B)** Timing of duplication of ParB. Schematic representation of ParB, PleC and DivJ localization during the cell cycle (left). Cells harboring a ParB-GFP fusion were tracked over time, and the first appearance of a second ParB protein was recorded, from the time of birth of each cell (right). Around 100 cells were analyzed, in three independent replicates. Statistical comparisons are calculated using unpaired *t*-tests (ns: not significant)**. (C)** Number of cells with a 1 or 2 ParB foci at the population level. Cells grown to mid exponential phase (OD600 = 0.4-0.6) were imaged by fluorescence microscopy, and the number of cells with 1, 2 or more ParB foci par field of view were calculated. Only cells exhibiting a fluorescent ParB signal are reported. Quantification for WT and Δ*hrmA* are shown in white and red respectively. Statistical comparisons are calculated using unpaired *t*-tests (ns: not significant)**. (D)** Number of cells with a localized PleC protein (indicative of a swarmer cell) over time. Synchronized cells were imaged by fluorescence microscopy and the number of cells per field of view with a localized PleC-YFP focus were quantified. The points are the average of 10 random images (around 100 cells) and three independent replicates for each time point, and the error bars represent the SEM. Data for WT and Δ*hmrA* are shown in black squares and red triangles respectively. **(E)** Number of cells with a localized DivJ protein (indicative of a stalked cell) over time. Synchronized cells were imaged by fluorescence microscopy and the number of cells per field of view with a localized DivJ-CFP focus were quantified. The points are the average of 10 random images (around 100 cells) and three independent replicates for each time point, and the error bars represent the SEM. Data for WT and Δ*hrmA* are shown in black squares and red triangles respectively.

**Figure S6:**
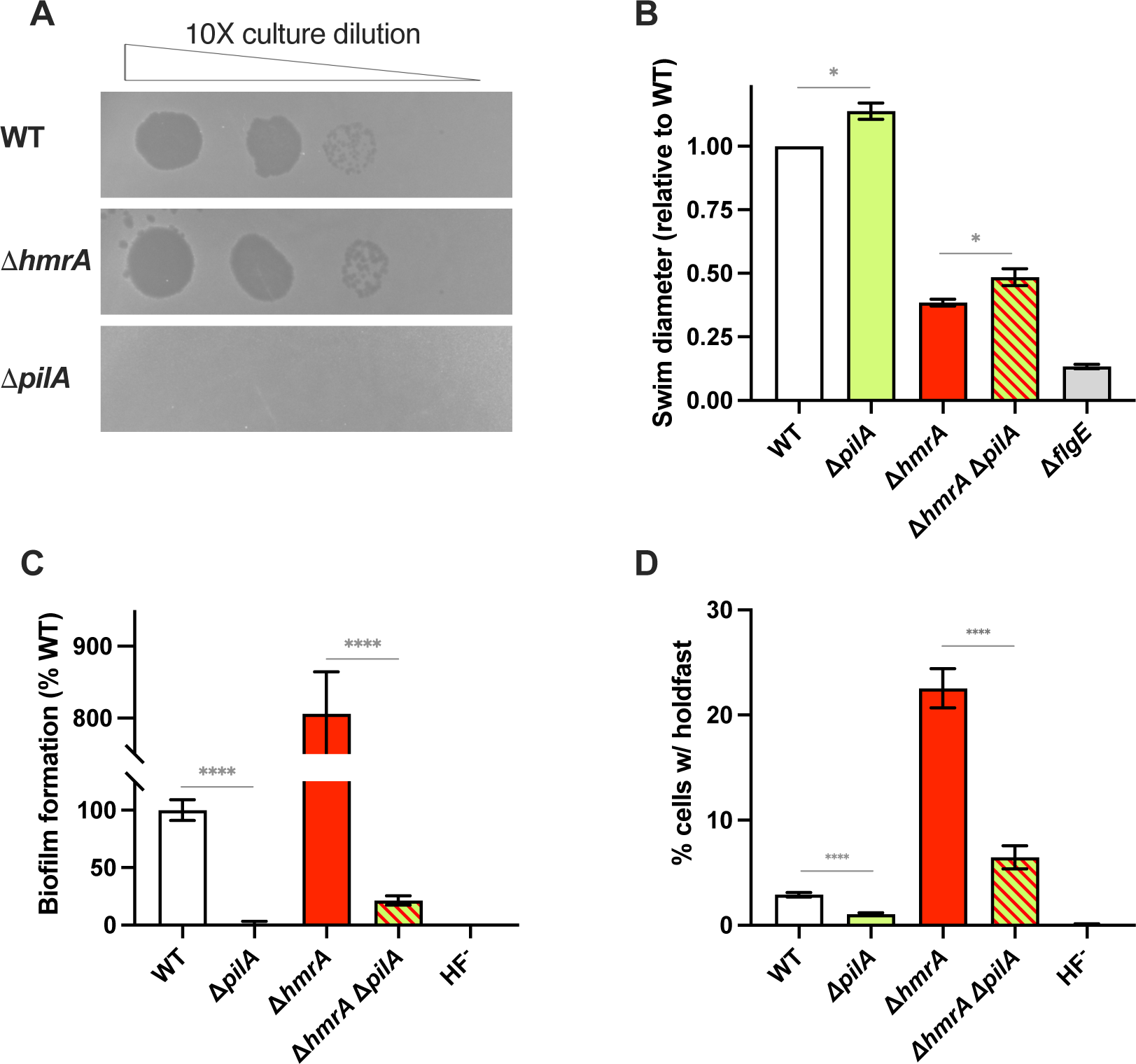
pili are not linked to the HmrA regulation pathway. **(A)** ΦCbK phage sensitivity assays. Cells were diluted to 10^-8^ cells per ml. Ten μl of the dilutions were spotted onto a PYE agar plate preabsorbed with ΦCbK phage. The plate was imaged after 24-hour incubation. **(B)** Motility assays in semisolid agar. Results are normalized to WT ring diameter measured on the same plate (set to 1). Bar graphs indicate the mean of three independent replicates, and error bars represent SEM. **(C)** Biofilm formation. Results are expressed as a percentage of biofilm formed by each strain compared to WT set at 100%. Results are given as the average of three independent experiments, each run in triplicate, and the error bars represent the SEM. **(D)** Quantification of cells harboring a holdfast in mixed populations. The results represent the average of at least three independent replicates (more than 300 cells per replicate) and the error bars represent the SEM. For all the graphs, statistical comparisons are calculated using unpaired *t*-tests. (* *P* < 0.05; ** *P* < 0.01; *** *P* < 0.001; **** *P* < 0.0001).

**Figure S7:**
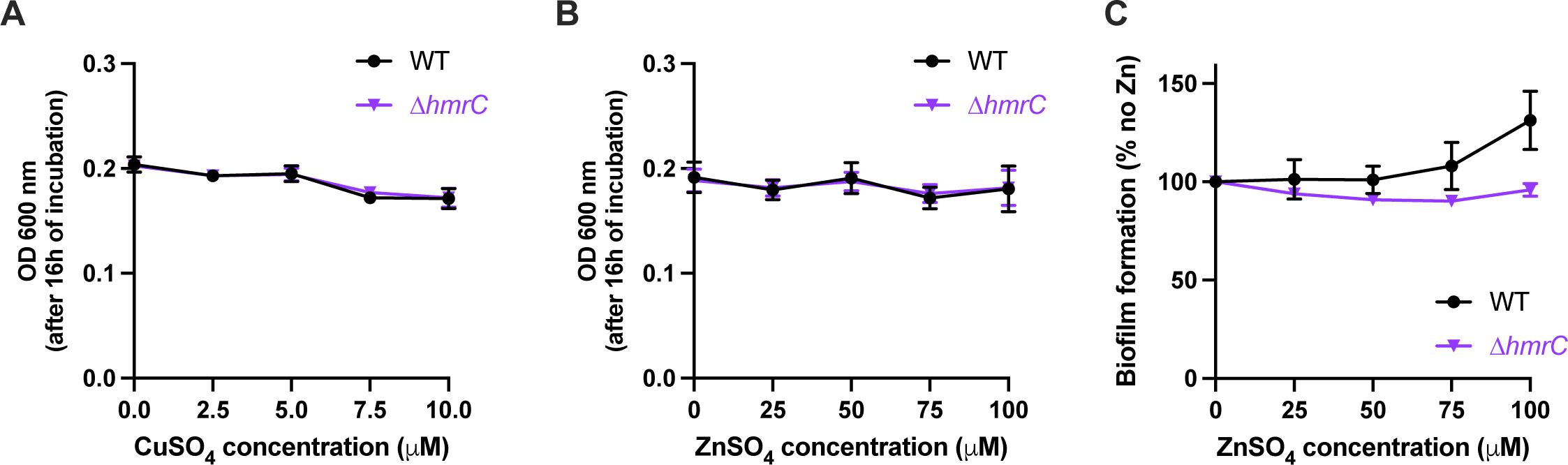
Growth and biofilm formation of WT and Δ*hmrC* in the presence of different metal concentrations. Growth after 24 h of incubation at 30°C, as measured by recording the absorbance at 600 nm (OD 600 nm) in the well of the 24-well plate used for biofilm assays, in the presence of CuSO_4_ **(A)** or ZnSO_4_ **(B)**. WT and Δ*hmrC* are shown in black circles and purple triangles respectively. **(C)** Biofilm formation in M2X medium adding different concentration of ZnSO_4_. Results are expressed as a percentage of biofilm formed by each strain in the presence of a set concentration of ZnSO_4_ compared to the no metal addition set at 100%. Results are given as at least five independent replicates, shown as black circles and purple triangles for WT and Δ*hmrC* respectively.

**Table S1:**
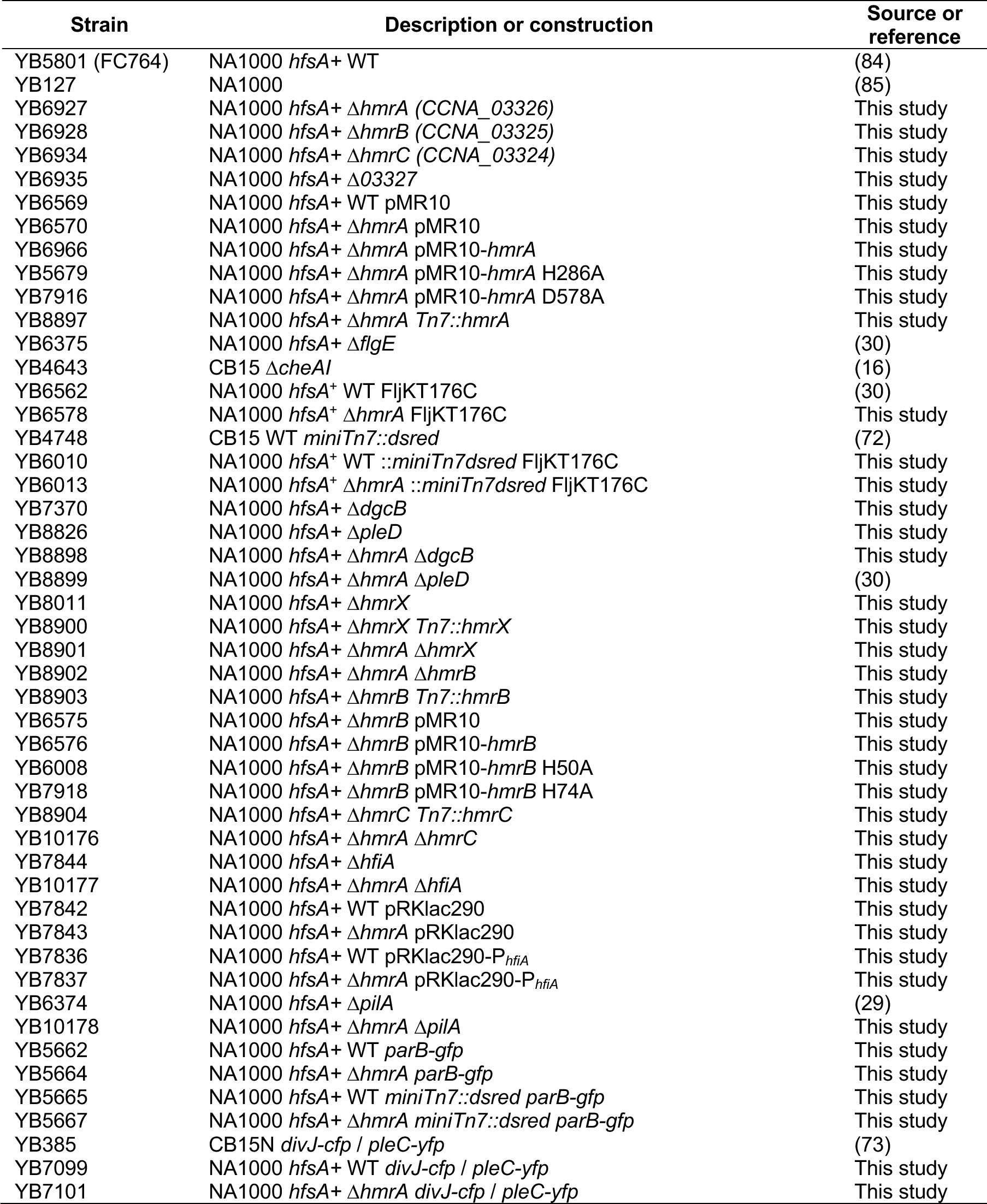
strains used in this study.

**Table S2:** plasmids used in this study.

| Plasmid | Description | Antibiotic | Reference |
| --- | --- | --- | --- |
| pNPTS138 | Litmus 38 derivative used for unmarked deletions in *C. crescentus* (*sacB* counter-selection) | Kan | (36) |
| pNPTS139 | Litmus 39 derivative used for unmarked deletions in *C. crescentus* (*sacB* counter-selection) | Kan | (36) |
| pNPTShmrA | pNPTS138 containing around 500 bp fragments upstream and downstream of *hmrA* (for in-frame deletion) | Kan | This study |
| pNPTShmrB | pNPTS138 containing around 500 bp fragments upstream and downstream of *hmrB* (for in-frame deletion) | Kan | This study |
| pNPTShmrC | pNPTS138 containing around 500 bp fragments upstream and downstream of *hmrC* (for in-frame deletion) | Kan | This study |
| pNPTShmrX | pNPTS138 containing around 500 bp fragments upstream and downstream of *hmrX* (for in-frame deletion) | Kan | This study |
| pNPTSdgcB | pNPTS138 containing around 500 bp fragments upstream and downstream of *dgcB* (for in-frame deletion) | Kan | This study |
| pNPTShfiA | pNPTS138 derivative used for in-frame deletion of *hfiA* |  | (15) |
| pNPTSfljKT176C | pNPTS139 containing 753-bp of *fljK* with a T to C point mutation at 103 bp from the start codon | Kan | (30) |
| pRKlac290 | plasmid for *lacZ* transcriptional fusions | Tc | (86) |
| pAF427 | pRKlac290-*PhfiA* | Tc | (15) |
| pMR10 | Mid copy replicating plasmid, IPTG inducible (constitutive in *C. crescentus*) | Kan | (87) |
| pMR10-*hmrA* | constitutive expression of *hmrA* | Kan | This study |
| pMR10-*hmrA* H286A | constitutive expression of *hmrAH286A* | Kan | This study |
| pMR10-*hmrA* D578A | constitutive expression of *hmrAD578A* | Kan | This study |
| pMR10-*hmrB* | constitutive expression of *hmrB* | Kan | This study |
| pMR10-*hmrB* H50A | constitutive expression of *hmrBH50A* | Kan | This study |
| pMR10-*hmrB* H74A | constitutive expression of *hmrBH74A* | Kan | This study |
| pMR10-*hmrC* | constitutive expression of *hmrC* | Kan | This study |
| pUC18-mini-Tn7T-*lac* | mini-Tn7 vector for insertion at the Tn7 *att* site | Gm | (69) |
| pTNS3 | helper plasmid for mating pTn7 in *C. crescentus* | Ap | (70) |
| pUC18-mini-Tn7T-*hmrA* | for *hmrA* insertion (native promoter) at the Tn7 *att* site | Gm | This study |
| pUC18-mini-Tn7T-*hmrB* | for *hmrB* insertion (native promoter) at the Tn7 *att* site | Gm | This study |
| pUC18-mini-Tn7T-*hmrC* | for *hmrC* insertion (native promoter) at the Tn7 *att* site | Gm | This study |
| pUC18-mini-Tn7T-*hmrX* | for *hmrX* insertion (native promoter) at the Tn7 *att* site | Gm | This study |
| pFD1 | *Mariner*-based mini-transposon vector for random transposon-mediated mutagenesis | Kan | (22) |

## REFERENCES

1. Amann RI, Ludwig W, Schleifer K-H. 1995. Phylogenetic identification and in situ detection of individual microbial cells without cultivation. Microbiol :143–169.

2. Costerton JW. 1995. Overview of microbial biofilms. J Ind Microbiol Biotechnol 15:137–140.

3. Davey ME, O’toole GA. 2000. Microbial biofilms: from ecology to molecular genetics. Microbiol Mol Biol Rev 64:847–867.

4. Hall-Stoodley L, Costerton JW, Stoodley P. 2004. Bacterial biofilms: from the natural environment to infectious diseases. Nat Rev Microbiol 2:95–108.

5. Fong JN, Yildiz FH. 2015. Biofilm matrix proteins. Microbiol Spectr 3:3.2. 28.

6. Berne C, Ducret A, Hardy GG, Brun YV. 2015. Adhesins involved in attachment to abiotic surfaces by Gram-negative bacteria. Microbiol Spectr 3.

7. Berne C, Ellison CK, Ducret A, Brun YV. 2018. Bacterial adhesion at the single-cell level. Nat Rev Microbiol 16:616–27.

8. Janakiraman RS, Brun YV. 1999. Cell cycle control of a holdfast attachment gene in *Caulobacter crescentus*. J Bacteriol 181:1118–1125.

9. Schrader JM, Li G-W, Childers WS, Perez AM, Weissman JS, Shapiro L, McAdams HH. 2016. Dynamic translation regulation in *Caulobacter* cell cycle control. Proc Natl Acad Sci USA 113:E6859–E6867.

10. Jenal U, Reinders A, Lori C. 2017. Cyclic di-GMP: second messenger extraordinaire. Nat Rev Microbiol 15:271–284.

11. Aldridge P, Paul R, Goymer P, Rainey P, Jenal U. 2003. Role of the GGDEF regulator PleD in polar development of *Caulobacter crescentus*. Mol Microbiol 47:1695–1708.

12. Levi A, Jenal U. 2006. Holdfast formation in motile swarmer cells optimizes surface attachment during *Caulobacter crescentus* development. J Bacteriol 188:5315–5318.

13. Abel S, Chien P, Wassmann P, Schirmer T, Kaever V, Laub MT, Baker TA, Jenal U. 2011. Regulatory cohesion of cell cycle and cell differentiation through interlinked phosphorylation and second messenger networks. Mol Cell 43:550–560.

14. Purcell EB, Siegal-Gaskins D, Rawling DC, Fiebig A, Crosson S. 2007. A photosensory two-component system regulates bacterial cell attachment. Proc Natl Acad Sci USA 104:18241–18246.

15. Fiebig A, Herrou J, Fumeaux C, Radhakrishnan SK, Viollier PH, Crosson S. 2014. A cell cycle and nutritional checkpoint controlling bacterial surface adhesion. PLoS Genet 10:e1004101.

16. Berne C, Brun YV. 2019. The two chemotaxis clusters in *Caulobacter crescentus* play different roles in chemotaxis and biofilm regulation. J Bacteriol 201:e00071–19.

17. Lori C, Kaczmarczyk A, de Jong I, Jenal U. 2018. A single-domain response regulator functions as an integrating hub to coordinate general stress response and development in alphaproteobacteria. mBio 9:e00809–18.

18. Eaton DS, Crosson S, Fiebig A. 2016. Proper control of *Caulobacter crescentus* cell surface adhesion requires the general protein chaperone DnaK. J Bacteriol 198:2631–2642.

19. Berne C. 2023. Sticky decisions: The multilayered regulation of adhesin production by bacteria. PLoS Genet 19:e1010648.

20. Wuichet K, Cantwell BJ, Zhulin IB. 2010. Evolution and phyletic distribution of two-component signal transduction systems. Curr Opin Microbiol 13:219–225.

21. Gao R, Mack TR, Stock AM. 2007. Bacterial response regulators: versatile regulatory strategies from common domains. Trends Biochem Sci 32:225–234.

22. Rubin EJ, Akerley BJ, Novik VN, Lampe DJ, Husson RN, Mekalanos JJ. 1999. *In vivo* transposition of mariner-based elements in enteric bacteria and mycobacteria. Proc Natl Acad Sci USA 96:1645–1650.

23. Geer LY, Domrachev M, Lipman DJ, Bryant SH. 2002. CDART: protein homology by domain architecture. Genome Res 12:1619–1623.

24. Artimo P, Jonnalagedda M, Arnold K, Baratin D, Csardi G, De Castro E, Duvaud S, Flegel V, Fortier A, Gasteiger E. 2012. ExPASy: SIB bioinformatics resource portal. Nucleic Acids Res 40:W597–W603.

25. Jacob-Dubuisson F, Mechaly A, Betton J-M, Antoine R. 2018. Structural insights into the signalling mechanisms of two-component systems. Nat Rev Microbiol 16:585–593.

26. Gutierrez S, Tyczynski WG, Boomsma W, Teufel F, Winther O. 2022. MembraneFold: Visualising transmembrane protein structure and topology. bioRxiv:2022.12. 06.518085.

27. Gumerov VM, Ortega DR, Adebali O, Ulrich LE, Zhulin IB. 2020. MiST 3.0: an updated microbial signal transduction database with an emphasis on chemosensory systems. Nucleic Acids Res 48:D459–D464.

28. Biondi EG, Skerker JM, Arif M, Prasol MS, Perchuk BS, Laub MT. 2006. A phosphorelay system controls stalk biogenesis during cell cycle progression in *Caulobacter crescentus*. Mol Microbiol 59:386–401.

29. Ellison CK, Kan J, Dillard RS, Kysela DT, Ducret A, Berne C, Hampton CM, Ke Z, Wright ER, Biais N. 2017. Obstruction of pilus retraction stimulates bacterial surface sensing. Science 358:535–538.

30. Berne C, Ellison CK, Agarwal R, Severin GB, Fiebig A, Morton III RI, Waters CM, Brun YV. 2018. Feedback regulation of *Caulobacter crescentus* holdfast synthesis by flagellum assembly via the holdfast inhibitor HfiA. Mol Microbiol 110:219–238.

31. McLaughlin M, Hershey DM, Reyes Ruiz LM, Fiebig A, Crosson S. 2022. A cryptic transcription factor regulates *Caulobacter* adhesin development. PLoS Genet 18:e1010481.

32. Reyes Ruiz LM, Fiebig A, Crosson S. 2019. Regulation of bacterial surface attachment by a network of sensory transduction proteins. PLoS Genet 15:e1008022.

33. Hershey DM, Fiebig A, Crosson S. 2019. A genome-wide analysis of adhesion in *Caulobacter crescentus* identifies new regulatory and biosynthetic components for holdfast assembly. mBio 10:e02273–18.

34. Del Medico L, Cerletti D, Schächle P, Christen M, Christen B. 2020. The type IV pilin PilA couples surface attachment and cell-cycle initiation in *Caulobacter crescentus*. Proc Natl Acad Sci USA 117:9546–9553.

35. Snyder RA, Ellison CK, Severin GB, Whitfield GB, Waters CM, Brun YV. 2020. Surface sensing stimulates cellular differentiation in *Caulobacter crescentus*. Proc Natl Acad Sci USA 117:17984–17991.

36. Alley M, Gomes SL, Alexander W, Shapiro L. 1991. Genetic analysis of a temporally transcribed chemotaxis gene cluster in *Caulobacter crescentus*. Genetics 129:333–341.

37. Wolfe AJ, Berg HC. 1989. Migration of bacteria in semisolid agar. Proc Natl Acad Sci USA 86:6973–6977.

38. Nesper J, Hug I, Kato S, Hee C-S, Habazettl JM, Manfredi P, Grzesiek S, Schirmer T, Emonet T, Jenal U. 2017. Cyclic di-GMP differentially tunes a bacterial flagellar motor through a novel class of CheY-like regulators. Elife 6:e28842.

39. Briegel A, Beeby M, Thanbichler M, Jensen GJ. 2011. Activated chemoreceptor arrays remain intact and hexagonally packed. Mol Microbiol 82:748–757.

40. Kovarik ML, Brown PJ, Kysela DT, Berne C, Kinsella AC, Brun YV, Jacobson SC. 2010. Microchannel-nanopore device for bacterial chemotaxis assays. Anal Chem 82:9357–9364.

41. Briegel A, Ding HJ, Li Z, Werner J, Gitai Z, Dias DP, Jensen RB, Jensen GJ. 2008. Location and architecture of the *Caulobacter crescentus* chemoreceptor array. Mol Microbiol 69:30–41.

42. Hershey DM. 2021. Integrated control of surface adaptation by the bacterial flagellum. Curr Opin Microbiol 61:1–7.

43. Hug I, Deshpande S, Sprecher KS, Pfohl T, Jenal U. 2017. Second messenger–mediated tactile response by a bacterial rotary motor. Science 358:531–534.

44. Holm L. 2022. Dali server: structural unification of protein families. Nucleic Acids Res 50:W210–W215.

45. Biondi EG, Reisinger SJ, Skerker JM, Arif M, Perchuk BS, Ryan KR, Laub MT. 2006. Regulation of the bacterial cell cycle by an integrated genetic circuit. Nature 444:899–904.

46. Surujon D, Ratner DI. 2016. Use of a probabilistic motif search to identify histidine phosphotransfer domain-containing proteins. PLoS One 11:e0146577.

47. Yang J, Yan R, Roy A, Xu D, Poisson J, Zhang Y. 2015. The I-TASSER Suite: protein structure and function prediction. Nat Meth 12:7–8.

48. Rempel S, Stanek W, Slotboom D. 2019. Energy-coupling factor-type ATP-binding cassette transporters. Annu Rev Biochem 88:551–576.

49. Park DM, Overton KW, Liou MJ, Jiao Y. 2017. Identification of a U/Zn/Cu responsive global regulatory two-component system in *Caulobacter crescentus*. Mol Microbiol 104:46–64.

50. de Araújo HL, Martins BP, Vicente AM, Lorenzetti AP, Koide T, Marques MV. 2021. Cold Regulation of genes encoding ion transport systems in the oligotrophic bacterium *Caulobacter crescentus*. Microbiol Spectr 9:e00710–21.

51. Skerker JM, Prasol MS, Perchuk BS, Biondi EG, Laub MT. 2005. Two-component signal transduction pathways regulating growth and cell cycle progression in a bacterium: a system-level analysis. PLoS Biol 3:e334.

52. Friedlander T, Mayo AE, Tlusty T, Alon U. 2015. Evolution of bow-tie architectures in biology. PLoS Comput Biol 11:e1004055.

53. Foreman R, Fiebig A, Crosson S. 2012. The LovK-LovR two-component system is a regulator of the general stress pathway in *Caulobacter crescentus*. J Bacteriol 194:3038–3049.

54. Ryder C, Byrd M, Wozniak DJ. 2007. Role of polysaccharides in *Pseudomonas aeruginosa* biofilm development. Curr Opin Microbiol 10:644–648.

55. Chambonnier G, Roux L, Redelberger D, Fadel F, Filloux A, Sivaneson M, De Bentzmann S, Bordi C. 2016. The hybrid histidine kinase LadS forms a multicomponent signal transduction system with the GacS/GacA two-component system in *Pseudomonas aeruginosa*. PLoS Genet 12:e1006032.

56. Willett JW, Crosson S. 2017. Atypical modes of bacterial histidine kinase signaling. Mol Microbiol 103:197–202.

57. Valentini M, Gonzalez D, Mavridou DA, Filloux A. 2018. Lifestyle transitions and adaptive pathogenesis of *Pseudomonas aeruginosa*. Curr Opin Microbiol 41:15–20.

58. Lourenço RF, Kohler C, Gomes SL. 2011. A two-component system, an anti-sigma factor and two paralogous ECF sigma factors are involved in the control of general stress response in *Caulobacter crescentus*. Mol Microbiol 80:1598–1612.

59. Fioravanti A, Fumeaux C, Mohapatra SS, Bompard C, Brilli M, Frandi A, Castric V, Villeret V, Viollier PH, Biondi EG. 2013. DNA binding of the cell cycle transcriptional regulator GcrA depends on N6-adenosine methylation in *Caulobacter crescentus* and other Alphaproteobacteria. PLoS Genet 9:e1003541.

60. Römling U, Galperin MY, Gomelsky M. 2013. Cyclic di-GMP: the first 25 years of a universal bacterial second messenger. Microbiol Mol Biol Rev 77:1–52.

61. Rood KL, Clark NE, Stoddard PR, Garman SC, Chien P. 2012. Adaptor-dependent degradation of a cell-cycle regulator uses a unique substrate architecture. Structure 20:1223–1232.

62. Maertens L, Cherry P, Tilquin F, Van Houdt R, Matroule J-Y. 2021. Environmental conditions modulate the transcriptomic response of both *Caulobacter crescentus* morphotypes to Cu stress. Microorganisms 9:1116.

63. Lawarée E, Gillet S, Louis G, Tilquin F, Le Blastier S, Cambier P, Matroule J-Y. 2016. *Caulobacter crescentus* intrinsic dimorphism provides a prompt bimodal response to copper stress. Nat Microbiol 1:1–7.

64. Shivaji S, Prakash JS. 2010. How do bacteria sense and respond to low temperature? Arch Microbiol 192:85–95.

65. Christen B, Abeliuk E, Collier JM, Kalogeraki VS, Passarelli B, Coller JA, Fero MJ, McAdams HH, Shapiro L. 2011. The essential genome of a bacterium.Mol Syst Biol 7:528.

66. Johnson RC, Ely B. 1977. Isolation of spontaneously derived mutants of *Caulobacter crescentus*. Genetics 86:25–32.

67. Poindexter JS. 1964. Biological properties and classification of the *Caulobacter* group. Bacteriol Rev 28:231.

68. Ried JL, Collmer A. 1987. An *nptI-sacB-sacR* cartridge for constructing directed, unmarked mutations in gram-negative bacteria by marker exchange-eviction mutagenesis. Gene 57:239–246.

69. Choi K-H, Gaynor JB, White KG, Lopez C, Bosio CM, Karkhoff-Schweizer RR, Schweizer HP. 2005. A *Tn7*-based broad-range bacterial cloning and expression system. Nat Meth 2:443–448.

70. Figueroa-Cuilan W, Daniel JJ, Howell M, Sulaiman A, Brown PJ. 2016. Mini-Tn7 insertion in an artificial *att* Tn7 site enables depletion of the essential master regulator CtrA in the phytopathogen *Agrobacterium tumefaciens*. Appl Environ Microbiol 82:5015–5025.

71. West L, Yang D, Stephens C. 2002. Use of the *Caulobacter crescentus* genome sequence to develop a method for systematic genetic mapping. J Bacteriol 184:2155–2166.

72. Berne C, Kysela DT, Brun YV. 2010. A bacterial extracellular DNA inhibits settling of motile progeny cells within a biofilm. Mol Microbiol 77:815–829.

73. Wheeler RT, Shapiro L. 1999. Differential localization of two histidine kinases controlling bacterial cell differentiation. Mol Cell 4:683–694.

74. Merker RI, Smit J. 1988. Characterization of the adhesive holdfast of marine and freshwater caulobacters. Appl Environ Microbiol 54:2078–2085.

75. Miller J. 1972. Experiment 48: Assay of β-galactosidase. Experiments in Molecular Genetics (Miller, JH, ed) pp. 352–355. Cold Spring Harbor Laboratory, Cold Spring Harbor, NY.

76. Schneider CA, Rasband WS, Eliceiri KW. 2012. NIH Image to ImageJ: 25 years of image analysis. Nat Meth 9:671.

77. Degnen ST, Newton A. 1972. Chromosome replication during development in *Caulobacter crescentus*. J Mol Biol 64:671–680.

78. Ducret A, Quardokus EM, Brun YV. 2016. MicrobeJ, a tool for high throughput bacterial cell detection and quantitative analysis. Nat Microbiol 1:1–7.

79. Nierman WC, Feldblyum TV, Laub MT, Paulsen IT, Nelson KE, Eisen J, Heidelberg JF, Alley M, Ohta N, Maddock JR. 2001. Complete genome sequence of *Caulobacter crescentus*. Proc Natl Acad Sci USA 98:4136–4141.

80. Phogat SK, Gupta R, Burma PK, Sen K, Pental D. 2001. On the estimation of number of events required for saturation mutagenesis of large genomes. Curr Sci 80:823–824.

81. Fernandez-Fernandez C, Gonzalez D, Collier J. 2011. Regulation of the activity of the dual-function DnaA protein in *Caulobacter crescentus*. PLoS One 6:e26028.

82. Pfaffl MW. 2001. A new mathematical model for relative quantification in real-time RT– PCR. Nucleic Acids Res 29:e45–e45.

83. Thanbichler M, Shapiro L. 2006. MipZ, a spatial regulator coordinating chromosome segregation with cell division in *Caulobacter*. Cell 126:147–162.

84. Marks ME, Castro-Rojas CM, Teiling C, Du L, Kapatral V, Walunas TL, Crosson S. 2010. The genetic basis of laboratory adaptation in *Caulobacter crescentus*. J Bacteriol 192:3678–3688.

85. Evinger M, Agabian N. 1977. Envelope-associated nucleoid from *Caulobacter crescentus* stalked and swarmer cells. J Bacteriol 132:294–301.

86. Gober JW, Shapiro L. 1992. A developmentally regulated *Caulobacter* flagellar promoter is activated by 3’enhancer and IHF binding elements. Mol Biol Cell 3:913–926.

87. Roberts RC, Toochinda C, Avedissian M, Baldini RL, Gomes SL, Shapiro L. 1996. Identification of a *Caulobacter crescentus* operon encoding *hrcA*, involved in negatively regulating heat-inducible transcription, and the chaperone gene *grpE*. J Bacteriol 178:1829–1841.

